# Comprehensive functional mapping of accessory chromosomes identifies a dominant virulence-regulator paralog in tomato wilt pathogen

**DOI:** 10.64898/2026.09.08.750270

**Authors:** Masaya Yamazaki, Hiroki Saito, Shuta Asai, Takashi Kamakura, Ken Komatsu, Takayuki Arazoe, Tsutomu Arie

## Abstract

Accessory chromosomes (ACs) serve as flexible genomic compartments that facilitate rapid adaptive evolution in eukaryotic microbes. In the tomato wilt pathogen *Fusarium oxysporum* f. sp. *lycopersici*, ACs are essential for virulence and host specificity; however, their structural complexity and functional redundancy have hindered a systematic characterization of their distinct roles. Here, we established a CRISPR/Cas9-based chromosomal dissection platform to generate a comprehensive functional map of pathogenicity determinants within ACs. Using this platform, we successfully generated a library of 36 large deletion mutants across the approximately 10 Mb putative AC region, enabling a chromosome-scale functional characterization of these compartments. A systematic screen of this library identified seven discrete AC segments that are indispensable for full virulence toward tomato. High-resolution chromosomal dissection of one virulence-associated segment through iterative subdivision and targeted gene disruption revealed that *FTF1a-1*, a single member of the *Fusarium transcription factor 1* (*FTF1*) family that originated from the duplication of core-chromosomal virulence gene *FTF2* into the AC region, functions as a dominant regulator of virulence. Given that the *FTF1a-3* paralog contributes only marginally to virulence and the deletion of other paralogs does not markedly affect disease development, our findings demonstrate a functional hierarchy within the duplicated *FTF1* gene family. These results imply that neofunctionalization of virulence genes within plastic fungal genomes promotes hyper-virulence and drives host-specific adaptation in *F. oxysporum*.

---

The genome organization of eukaryotes varies widely across species. Notably, some eukaryotic microbes harbor accessory chromosomes (ACs) that are dispensable for basic survival but confer specialized adaptive traits under specific environmental conditions^1–4^. In plant-pathogenic fungi, these non-essential genomic compartments serve as reservoirs of virulence-related genes that drive host-specific adaptation during colonization. These include genes encoding host-specific toxins and other secondary metabolites, as well as secreted effector proteins that suppress host immunity^5–12^.

*Fusarium oxysporum* ranks among the top ten most economically and scientifically important plant-pathogenic fungi, causing devastating diseases in more than 100 plant species worldwide^13^. This fungus comprises a species complex of morphologically indistinguishable lineages, many of which exhibit host-specific pathogenicity. As a model for investigating the mechanisms of pathogenicity and host specificity, *F. oxysporum* f. sp. *lycopersici* (Fol), the causal agent of tomato (*Solanum lycopersicum*) wilt, was the first *forma specialis* (f. sp.) within *F. oxysporum* to have its genome sequenced^5^. The reference genome of Fol isolate 4287 (race 2) consists of 15 chromosomes, four of which are ACs (chromosomes 3, 6, 14, and 15), as well as additional AC regions located at the ends of core chromosomes 1 and 2^5^. Remarkably, horizontal transfer of chromosome 14 alone into a nonpathogenic *F. oxysporum* isolate is sufficient to confer pathogenicity on tomato, providing direct experimental evidence that this mobile AC, designated the pathogenicity chromosome, dictates the host-specific pathogenicity of this fungus^5,14^.

Highly enriched in transposable elements, repetitive sequences, and AT-rich regions, AC compartments are substantially more prone to spontaneous loss and structural rearrangements than core chromosomes^4,14^. In *F. oxysporum*, this plasticity allows the ACs to gain and lose virulence-related genes that are specifically induced during infection, exemplified by the Secreted in xylem (Six) effectors^15–17^. To date, fourteen Six effectors have been identified in the xylem sap of Fol-infected tomato plants^5,15,18,19^. Secreted into the host xylem, these effectors promote infection partly through suppression of host immunity, and four of them (Six1, Six3, Six5, and Six6) are required for its full virulence^16,18,20–22^. Besides effectors, ACs also encode expanded families of virulence-associated transcription factors that induce effector gene expression during infection^23^. The repertoires of AC-encoded virulence determinants, including effectors and transcription factors, differ among *formae speciales* and are thought to determine host specificity^24^. However, a high-resolution functional map of the specific AC regions and genes governing the infection process has yet to be established due to the structural complexity of ACs and the functional redundancy of AC-encoded virulence genes. To address this gap, we optimized a CRISPR/Cas9-mediated chromosomal dissection platform for generating large-scale deletions in the ACs of Fol isolate MAFF 103036 (race 1) harboring the full complement of *SIX* effectors (Supplementary Table 1). Systematic characterization of the resulting AC deletion library, covering 99% of the targeted AC regions, revealed that multiple AC segments are required for full virulence and led to the identification of a dominant virulence regulator among duplicated paralogs.

## Results

### Chromosome-level genome assembly of Fol MAFF 103036

To enable the construction of a large-scale AC deletion library, we generated a chromosome-level genome assembly of Fol MAFF 103036 using the PacBio Sequel SMRT platform. The assembly yielded a high-quality genome (BUSCO score, 99.3%) comprising 19 chromosome-level contigs, including nine with telomeric repeats at both ends, nine with a single telomeric repeat at one end, and one lacking telomeric sequences (Table 1). RNA-seq-supported annotation identified 16,753 protein-coding genes in the assembled genome (Table 1). Comparative genomic analysis against the well-characterized reference genome of Fol 4287 revealed that a part of contig 5 and the entirety of contigs 6, 11, 14, 15, 16, 18, and 19 in MAFF 103036 are syntenic with the AC regions of 4287 (Fig. 1a and Supplementary Fig. 1a, d). These putative AC regions exhibited a high density of repetitive sequences and a low density of predicted genes (Supplementary Fig. 1b, c), consistent with the hallmark features of *F. oxysporum* AC regions. Notably, a part of contig 14 and the entire contigs 15 and 16 showed high synteny with the pathogenicity chromosome 14 of 4287 (Supplementary Fig. 1a, d).

**Figure 1.**
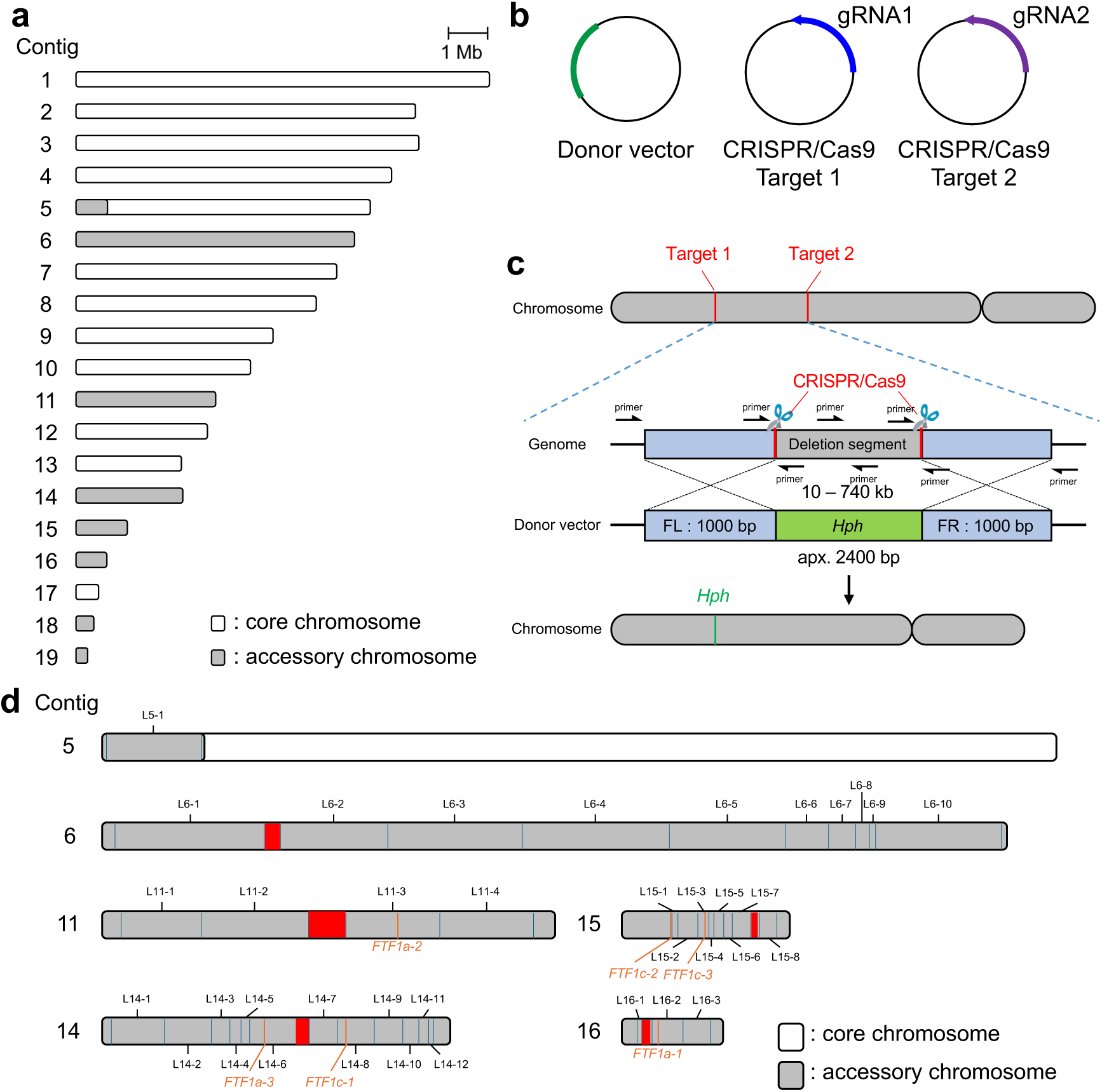
Construction of an AC deletion library in MAFF 103036 using CRISPR/Cas9-mediated genome editing. **a,** Genomic organization of ACs and core chromosomes in MAFF 103036. **b,** Schematic representation of vectors for protoplast transformation. One donor vector and two CRISPR/Cas9 vectors were co-transformed into protoplasts. **c,** Schematic representation of CRISPR/Cas9-mediated large-scale deletion of AC segments. Deletions were validated using primer sets designed outside the upstream and downstream homology arms. Additionally, three internal primer pairs were utilized to verify the excision of internal sequences. FL, flanking region on the left side; FR, flanking region on the right side; *Hph*, *hygromycin phosphotransferase* cassette. **d,** Structural organization of ACs in MAFF 103036. Segments targeted for large-scale deletion between CRISPR/Cas9 cleavage sites are designated by their segment names. Red boxes, AT-rich regions; blue bars, CRISPR/Cas9 target sites; orange bars, *FTF* paralogs.

**Table 1.** Genome assembly and annotation statistics for MAFF 103036.

| Analysis | Statistic | MAFF 103036 |
| --- | --- | --- |
| Assembly |  |  |
|  | Mean coverage (fold) | 195 |
|  | Assembly size (bp) | 57292497 |
|  | No. of contigs | 19 |
|  | Maximum contig length (bp) | 6570833 |
|  | N<sub>50</sub> contig length (bp) | 4427170 |
|  | GC content (%) | 47.81 |
|  | BUSCO* coverage (%) | 99.3 |
| Presence of telomeric repeats |  |  |
|  | On both ends | 9 |
|  | On one end | 9 |
|  | None | 1 |
| Predicted genes |  |  |
|  | Total no. of genes | 16753 |
|  | Total no. of proteins | 18410 |

741 \*Sordariomycetes\_odb12

### Development of a CRISPR/Cas9-mediated large-scale chromosomal deletion platform

The vector-based CRISPR/Cas9 system optimized for *F. oxysporum* enables efficient homologous recombination-mediated gene deletion within ACs^25^. We therefore hypothesized that introducing simultaneous DNA double-strand breaks at both ends of a targeted segment would promote homologous recombination with a donor vector harboring a *hygromycin phosphotransferase* (*Hph*) cassette, thereby enabling large-scale deletions of AC regions (Fig. 1b, c). To validate this strategy, we targeted a 59-kb segment on the predicted AC contig 15, designated L15-3 (Fig. 1d and Table 2). Two CRISPR/Cas9 vectors, each harboring one of the target sequences, were co-transformed into MAFF 103036 protoplasts together with the donor vector via PEG-mediated transformation (Fig. 1b, c). Across three independent experiments, we obtained a total of 63 hygromycin-resistant transformants and screened them by PCR using primer pairs flanking the homology arms (Fig. 1c and Supplementary Table 2). Among these, one transformant yielded the expected PCR products (Supplementary Fig. 2a), and subsequent sequencing confirmed that the *Hph* cassette had precisely replaced the target segment. To exclude the possibility that the deleted segment had ectopically reintegrated into other genomic loci, this transformant was further analyzed using primer pairs targeting three distinct internal loci within the deleted region (Supplementary Table 2). The absence of amplicons at all internal loci confirmed the successful deletion of the entire L15-3 segment (Supplementary Fig. 2b). This mutant was used for subsequent functional analyses.

**Table 2.**
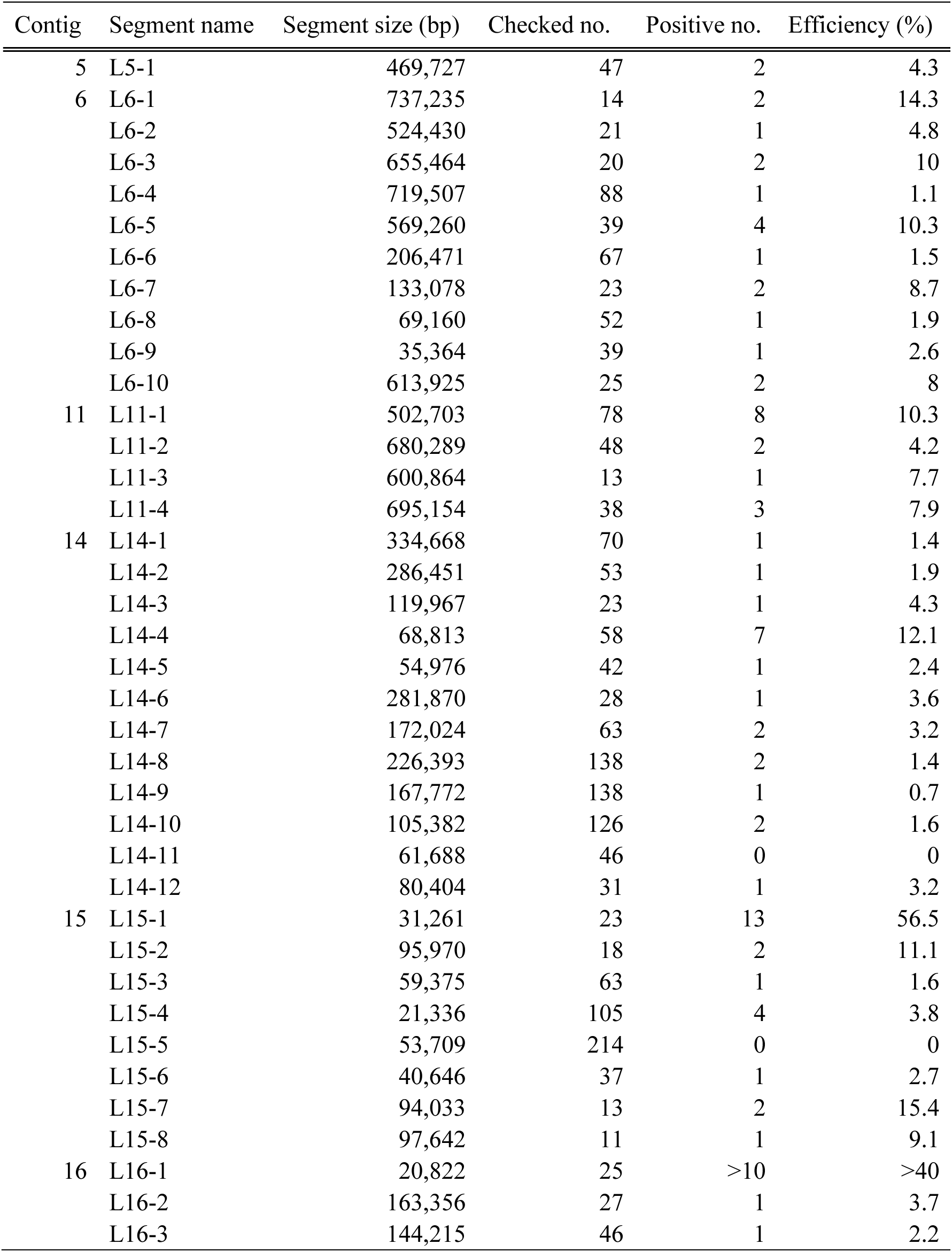
Construction efficiency of the AC deletion library in MAFF 103036.

| Contig | Segment name | Segment size (bp) | Checked no. | Positive no. | Efficiency (%) |
| --- | --- | --- | --- | --- | --- |
| 5 | L5-1 | 469,727 | 47 | 2 | 4.3 |
| 6 | L6-1 | 737,235 | 14 | 2 | 14.3 |
|  | L6-2 | 524,430 | 21 | 1 | 4.8 |
|  | L6-3 | 655,464 | 20 | 2 | 10 |
|  | L6-4 | 719,507 | 88 | 1 | 1.1 |
|  | L6-5 | 569,260 | 39 | 4 | 10.3 |
|  | L6-6 | 206,471 | 67 | 1 | 1.5 |
|  | L6-7 | 133,078 | 23 | 2 | 8.7 |
|  | L6-8 | 69,160 | 52 | 1 | 1.9 |
|  | L6-9 | 35,364 | 39 | 1 | 2.6 |
|  | L6-10 | 613,925 | 25 | 2 | 8 |
| 11 | L11-1 | 502,703 | 78 | 8 | 10.3 |
|  | L11-2 | 680,289 | 48 | 2 | 4.2 |
|  | L11-3 | 600,864 | 13 | 1 | 7.7 |
|  | L11-4 | 695,154 | 38 | 3 | 7.9 |
| 14 | L14-1 | 334,668 | 70 | 1 | 1.4 |
|  | L14-2 | 286,451 | 53 | 1 | 1.9 |
|  | L14-3 | 119,967 | 23 | 1 | 4.3 |
|  | L14-4 | 68,813 | 58 | 7 | 12.1 |
|  | L14-5 | 54,976 | 42 | 1 | 2.4 |
|  | L14-6 | 281,870 | 28 | 1 | 3.6 |
|  | L14-7 | 172,024 | 63 | 2 | 3.2 |
|  | L14-8 | 226,393 | 138 | 2 | 1.4 |
|  | L14-9 | 167,772 | 138 | 1 | 0.7 |
|  | L14-10 | 105,382 | 126 | 2 | 1.6 |
|  | L14-11 | 61,688 | 46 | 0 | 0 |
|  | L14-12 | 80,404 | 31 | 1 | 3.2 |
| 15 | L15-1 | 31,261 | 23 | 13 | 56.5 |
|  | L15-2 | 95,970 | 18 | 2 | 11.1 |
|  | L15-3 | 59,375 | 63 | 1 | 1.6 |
|  | L15-4 | 21,336 | 105 | 4 | 3.8 |
|  | L15-5 | 53,709 | 214 | 0 | 0 |
|  | L15-6 | 40,646 | 37 | 1 | 2.7 |
|  | L15-7 | 94,033 | 13 | 2 | 15.4 |
|  | L15-8 | 97,642 | 11 | 1 | 9.1 |
| 16 | L16-1 | 20,822 | 25 | >10 | >40 |
|  | L16-2 | 163,356 | 27 | 1 | 3.7 |
|  | L16-3 | 144,215 | 46 | 1 | 2.2 |

### Construction of an AC deletion library

Following the successful deletion of the L15-3 segment, we constructed a large-scale deletion library targeting the putative ACs (Fig. 1a). AT-rich regions (GC content <34%), which are putative centromeres, were excluded from targeting. Repeat-rich regions at contig ends were also excluded, as their repetitive sequences prevented the design of unique guide RNAs and homology arms. In addition, the small contigs 18 and 19 were excluded because they are likely terminal fragments of other ACs rather than independent chromosome-level contigs. Based on these criteria, we divided the putative AC regions into 38 segments ranging from 20 to 740 kb, comprising one on contig 5, ten on contig 6, four on contig 11, twelve on contig 14, eight on contig 15, and three on contig 16, for a total of approximately 10 Mb (Fig. 1d and Table 2). We successfully obtained deletion mutants for 36 of the 38 segments (Table 2 and Supplementary Figs. 2–9). These mutants collectively covered ∼9.9 Mb of the ∼10 Mb of AC regions targeted in this study (99% coverage; Table 2). Only one segment on contig 14 (L14-11) and one on contig 15 (L15-5) could not be deleted (Table 2). Deletion efficiency, defined as the percentage of hygromycin-resistant transformants carrying the intended deletion as verified by PCR, ranged from 0.7% to 56% and showed no apparent relationship with segment length (Table 2). This near-complete coverage enabled systematic functional analyses of the ACs.

### Seven discrete AC segments are required for full virulence on tomato

To identify AC segments contributing to virulence, we performed disease assays on tomato (cv. Ponderosa) using the 36 deletion mutants. Most mutants exhibited external symptoms comparable to those of the wild-type (Fig. 2a–f, h and Supplementary Fig. 10a–f, h). However, seven mutants (Δ L14-5, ΔL14-8, ΔL14-10, ΔL14-12, ΔL15-2, ΔL15-3, and ΔL16-2) showed significantly attenuated external symptoms (Fig. 2d–h and Supplementary Fig. 10d–h). Internal symptoms, assessed by the extent of vascular browning, showed consistent trends (Supplementary Fig. 11a–h). These results demonstrate that the seven discrete AC segments are required for full virulence of MAFF 103036. All mutants exhibited growth rates on PDA comparable to the wild-type (Supplementary Fig. 12), indicating that the reduced virulence was not due to general growth defects.

**Figure 2.**
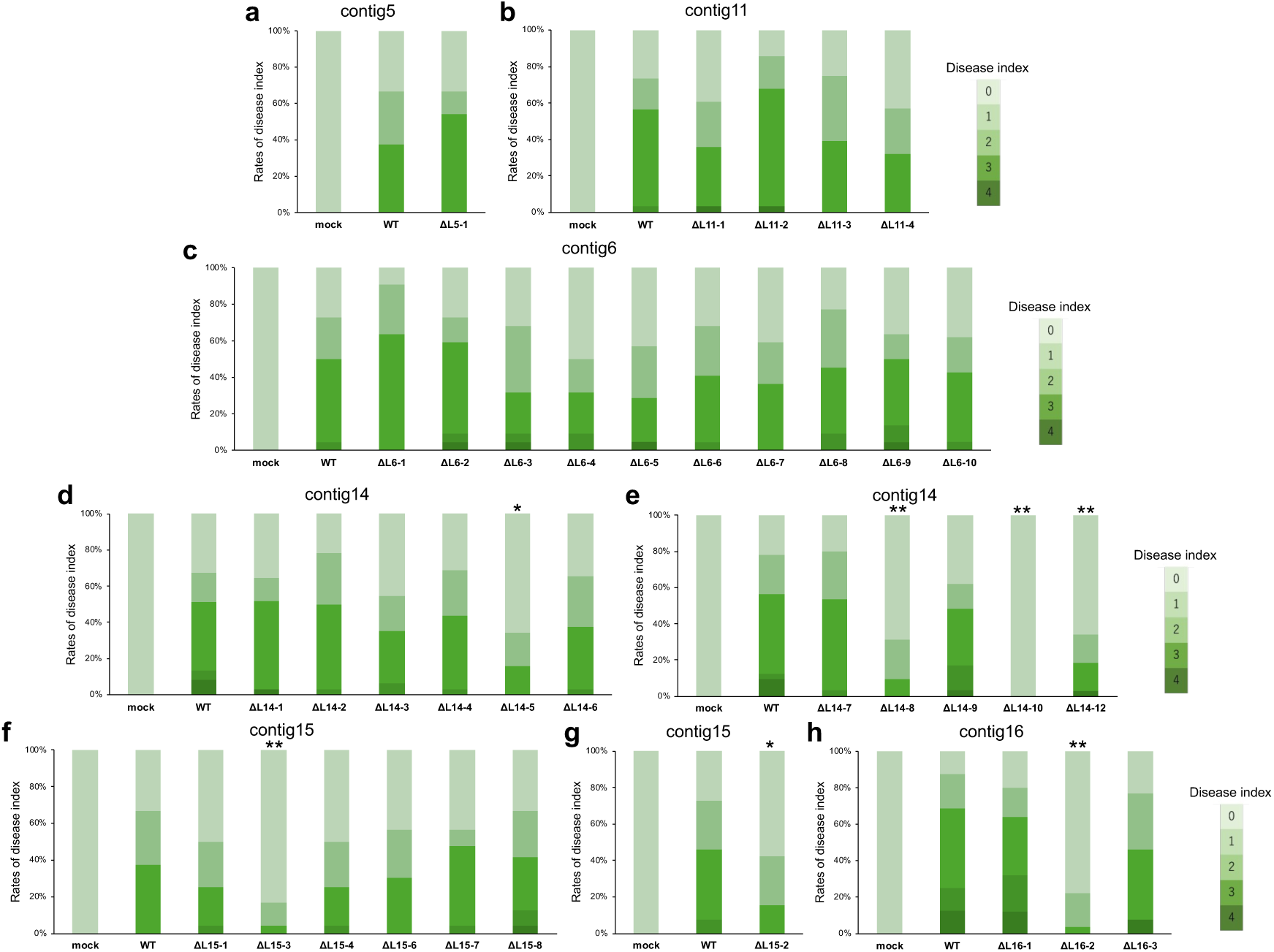
Virulence analysis of the AC deletion library in MAFF 103036 based on external disease symptoms. **a–h,** Deletion mutants corresponding to segments on contig 5 (a), contig 11 (b), contig 6 (c), the long arm of contig 14 (d), the short arm of contig 14 (e), contig 15 (f and g), and contig 16 (h) were evaluated for external disease symptoms. Disease index was scored as described in the Methods. Asterisks indicate significant differences compared with the wild-type (n = 21–37 biological replicates; *adjusted *P* < 0.05, **adjusted *P* < 0.01; Mann–Whitney *U*-test).

To identify candidate virulence genes, we compared expression profiles of the wild-type between *in vitro* and *in planta* conditions using RNA-seq (Supplementary Table 1). We analyzed the expression of genes in the seven segments identified above, as well as in L14-11 and L15-5, to analyze these two undeletable segments for candidate virulence genes as well. Previous studies have shown that several effector genes required for Fol virulence are located within the identified segments, including *SIX6* in L14-5 (Fig. 3a)^20^, *SIX5* and *SIX3* in L15-2 (Fig. 3f)^16,21,22^, and *SIX1* in L15-3 (Fig. 3g)^18^. The expression of these effectors was induced exclusively under *in planta* conditions (Fig. 3a, f, g), consistent with the reduced virulence of the corresponding deletion mutants. Therefore, our deletion library effectively captures virulence-related loci.

**Figure 3.**
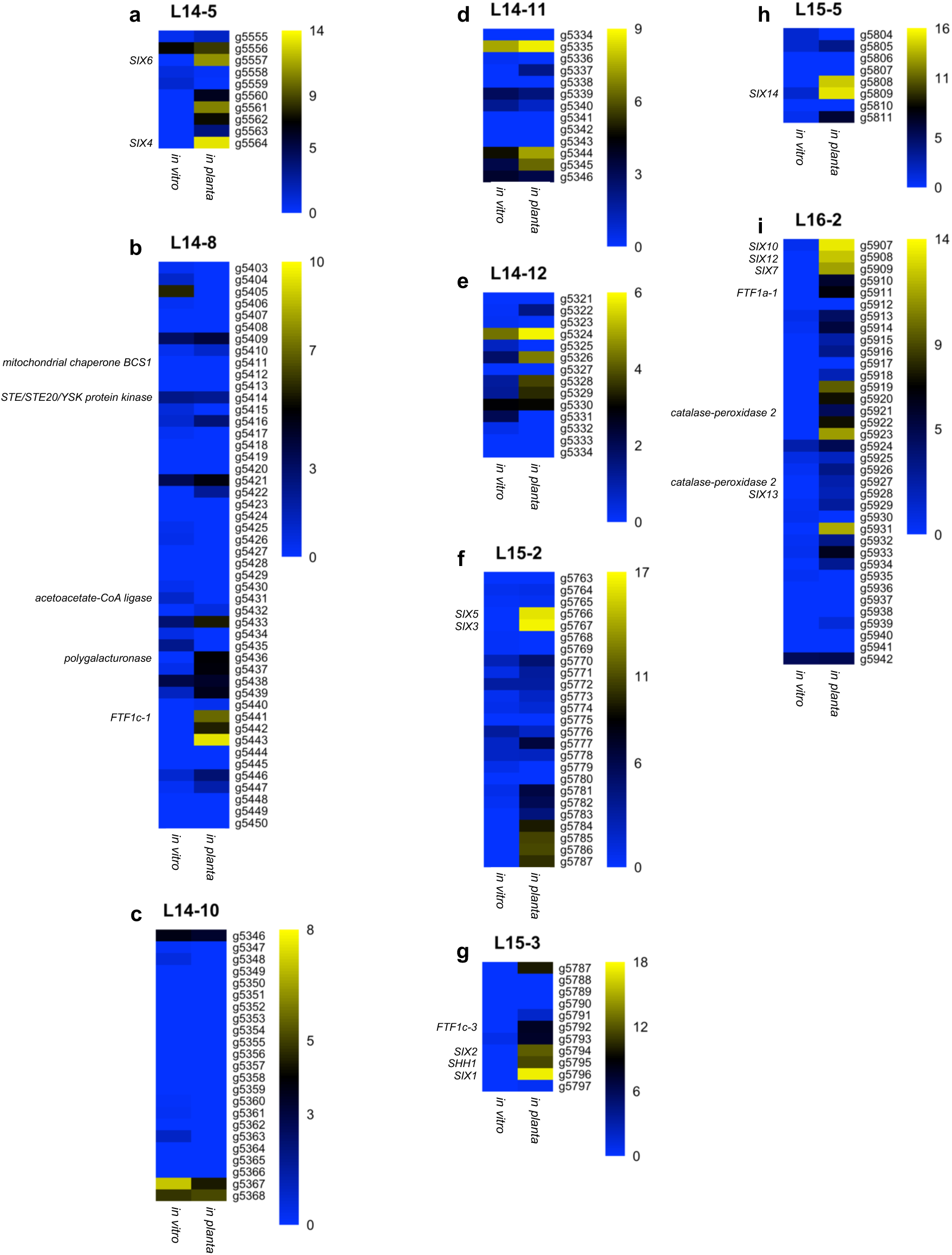
Transcriptome analysis of virulence-related segments identified in the AC deletion library and non-deleted segment of MAFF 103036. **a–i**, Heat maps showing the expression profiles of genes within the indicated segments (L14-5, L14-8, L14-10, L14-11, L14-12, L15-2, L15-3, L15-5, and L16-2) under *in vitro* and *in planta* conditions. Each row represents a gene, and each column represents a condition. Displayed values represent log₂-transformed TPM. The color scale indicates relative expression levels, with blue and yellow representing low and high expression, respectively. Gene names are indicated on the left side of each heatmap. Only genes exhibiting high sequence similarity to those of 4287 are included. Genes significantly upregulated under *in planta* conditions compared with *in vitro* conditions were defined as candidate virulence-related genes.

One of these seven segments, L16-2, contained the effectors *SIX10*, *SIX12*, and *SIX7*, as well as *FTF1a* (*Fol_MAFF103036_v2_g5911*), a member of the *Fusarium Transcription Factor 1* (*FTF1*) family implicated in effector regulation^26–28^. Consistent with previous reports, these genes were strongly upregulated under *in planta* conditions (Fig. 3i). Interestingly, the remaining segments (L14-8, L14-10, and L14-12), together with the two undeletable segments (L14-11 and L15-5), did not harbor any genes previously shown to be individually involved in virulence (Fig. 3b–e, h and Supplementary Table 3). Nevertheless, several genes within these segments were either upregulated or downregulated during plant colonization (Fig. 3b–e, h and Supplementary Table 3), suggesting the presence of novel virulence genes. Together, these findings demonstrate that our deletion library provides a systematic means of identifying virulence genes across the ACs.

### Stepwise dissection of the virulence-related segment L16-2

Given that L16-2 is a large segment spanning 163,356 bp and potentially encodes several unverified virulence-associated genes (Fig. 4a), we divided this region into three smaller subsegments (L16-2-1, L16-2-2, and L16-2-3) to assess their individual contributions to virulence (Fig. 4a and Supplementary Fig. 13a–f). Among these, only the ΔL16-2-1 mutant exhibited a clear reduction in virulence, displaying a phenotype comparable to that of the parental ΔL16-2 mutant (Fig. 4b and Supplementary Figs. 14a and 15a). The L16-2-1 segment spans 63,673 bp and contains 16 annotated genes. Four of these–*SIX10*, *SIX12*, *SIX7*, and *Fol_MAFF103036_v2_g5922*–are present as single-copy genes in the genome, whereas the remaining 12 are multicopy genes with high sequence similarity to loci elsewhere in the genome. Compared with multicopy genes, single-copy genes generally exhibit lower functional redundancy and are therefore more likely to be associated with essential or conserved functions^29,30^. Among the single-copy genes, *SIX10*, *SIX12*, and *SIX7* formed a 4,973-bp mini-cluster (Fig. 4a), and the amino acid sequence of *Fol_MAFF103036_v2_g5922* shows high similarity to that of a previously reported metalloproteinase (*MET*) gene located on the AC of the celery (*Apium graveolens*) yellows fungus *F. oxysporum* f. sp. *apii* (Fig. 4a)^31^. To determine whether these single-copy genes contribute to virulence, we generated individual deletion mutants and conducted disease assays (Supplementary Fig. 16a–d). Notably, none of these single-copy gene deletion mutants exhibited a significant difference in virulence compared with the wild-type (Supplementary Figs. 14b, c and 15b–e). These findings indicate that the determinants required for full virulence reside among the multicopy, rather than the single-copy, genes within L16-2-1. To pinpoint the specific determinants within this segment, we further subdivided L16-2-1 into three smaller subsegments (L16-2-1-1, L16-2-1-2 and L16-2-1-3) for functional characterization (Fig. 4a and Supplementary Fig. 17a–f). Only the ΔL16-2-1-1 mutant exhibited a significant reduction in virulence, fully recapitulating the phenotype observed in the parental ΔL16-2-1 mutant (Fig. 4c, d and Supplementary Figs. 14d, e and 15f, g). These results clearly indicate that the virulence-associated multicopy genes are localized within the L16-2-1-1 segment (Fig. 4a).

**Figure 4.**
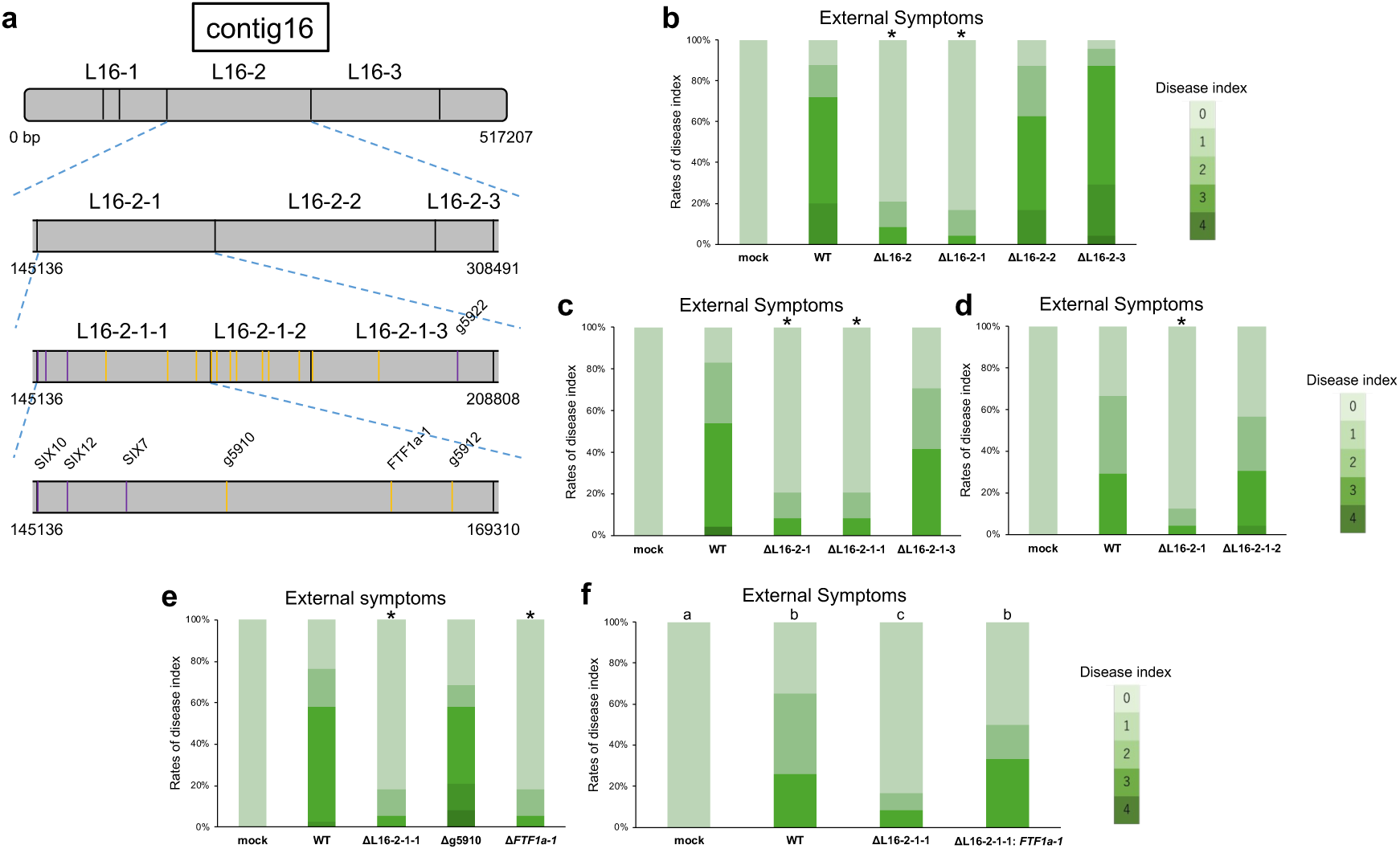
High-resolution deletion analysis of the virulence-associated L16-2 segment. **a**, Schematic representation of stepwise deletions within L16-2, a virulence-related segment on contig 16. Purple bars, single-copy genes; yellow bars, multicopy genes. **b–e**, External disease symptom-based evaluation of mutants generated by stepwise deletion of L16-2 and by the targeted disruption of genes within L16-2-1-1. The disease index was scored as described in the Methods. Asterisks indicate significant differences compared with the wild-type (n = 23–38 biological replicates; *adjusted *P* < 0.01; Mann–Whitney *U*-test). **f**, External symptom-based evaluation of a mutant strain in which the *FTF1a-1* locus was re-introduced into ΔL16-2-1-1 (ΔL16-2-1-1: *FTF1a-1*). Different letters indicate significant differences among the wild-type and mutant strains (n = 29–32 biological replicates; adjusted *P* < 0.05; Kruskal–Wallis test followed by Dunn’s multiple-comparison test).

### Transcription factor FTF1a-1 contributes to virulence

The L16-2-1-1 segment spans 24,175 bp and contains three multicopy genes, designated *Fol_MAFF103036_v2_g5910, FTF1a-1*, and *Fol_MAFF103036_v2_g5912* (Fig. 4a). *FTF1* family members were first characterized in the bean (*Phaseolus vulgaris*) pathogen *F. oxysporum* f. sp. *phaseoli*^26^ and are widely conserved among plant-pathogenic *F. oxysporum* isolates^27^. Although the broader role of *FTF1* in virulence has been investigated using gene-silencing approaches^27^, the specific contribution of each paralog remains elusive due to their high sequence similarity. To determine whether these genes contribute to virulence, we generated individual deletion mutants using the CRISPR/Cas9 system. Specific gRNAs were successfully designed for two of the three multicopy genes, *Fol_MAFF103036_v2_g5910* and *FTF1a-1*, enabling the construction of the corresponding deletion mutants (Supplementary Fig. 18a, b). Although the gRNA designed for *FTF1a-1* contained potential off-target sites in two paralogs, *Fol_MAFF103036_v2_g5441* and *Fol_MAFF103036_v2_g5754*, sequencing analysis of the resulting Δ*FTF1a-1* mutant confirmed that no unintended mutations were introduced at either predicted off-target site. Disease assays revealed that the Δ*FTF1a-1* mutant, but not the Δ*g5910* mutant, exhibited a significant reduction in virulence, displaying a phenotype comparable to that of the ΔL16-2-1-1 mutant (Fig. 4e and Supplementary Figs. 14f and 15h). Furthermore, ectopic complementation of the ΔL16-2-1-1 mutant with *FTF1a-1* under the control of its native promoter (ΔL16-2-1-1: *FTF1a-1*) restored virulence to wild-type levels (Fig. 4f, Supplementary Fig. 18c and Supplementary Figs. 14g and 15i), although *FTF1a-1* expression was not fully recovered to wild-type levels (Supplementary Fig. 19). This indicates that *FTF1a-1* is the primary virulence determinant within L16-2-1-1, and the remaining multicopy gene *Fol_MAFF103036_v2_g5912* is dispensable for virulence.

### FTF1a-1 is a primary virulence-associated paralog

To evaluate the functional contributions of the *FTF1* paralogs, we first examined the *FTF1* paralog repertoire in the MAFF 103036 genome. Ftf transcription factors are classified into two groups: Ftf1 and Ftf2. *FTF2* (*Fol_MAFF103036_v2_g16238*) exists as a single-copy gene on a core chromosome and is highly conserved across filamentous fungi^27,32^. In contrast, *FTF1* is subdivided into three distinct types—*FTF1a, FTF1b*, and *FTF1c*—based on the length of their coding sequences; notably, the promoter sequence of *FTF1c* also differs from those of *FTF1a* and *FTF1b*^27^. These *FTF1* genes are thought to have arisen as paralogs of *FTF2* during the evolution of pathogenicity^27^. In the MAFF 103036 genome, we identified three copies each of *FTF1a* and *FTF1c* located on the putative ACs. For clarity in subsequent analyses, we designated these additional copies as *FTF1a-2* (*Fol_MAFF103036_v2_g3393*), *FTF1a-3* (*Fol_MAFF103036_v2_g5534*), *FTF1c-1* (*Fol_MAFF103036_v2_g5441*), *FTF1c-2* (*Fol_MAFF103036_v2_g5754*), and *FTF1c-3* (*Fol_MAFF103036_v2_g5792*).

Chromosomal mapping revealed that *FTF1a-2*, *FTF1a-3*, *FTF1c-1*, and *FTF1c-3* are located on L11-3, L14-6, L14-8, and L15-3, respectively, whereas *FTF1c-2* is located outside the region covered by the AC deletion library (Fig. 1d). We generated five individual deletion mutants, Δ*FTF1a-2*, Δ*FTF1a-3*, Δ*FTF1c-1*, Δ*FTF1c-2*, and Δ*FTF1c-3* (Supplementary Fig. 20a–e), and assessed their virulence. Among the five mutants, only Δ*FTF1a-3* exhibited a significant reduction in virulence, whereas the others were indistinguishable from the wild-type (Supplementary Fig. 21a–i). Δ*FTF1a-1* exhibited a significantly stronger reduction in virulence than Δ*FTF1a-3* (Supplementary Fig. 21d, e), indicating that *FTF1a-1* and *FTF1a-3* both contribute to virulence on tomato, with *FTF1a-1* playing the predominant role among the analyzed paralogs (Supplementary Fig. 21a, d, e). Notably, Ftf1a-1 shares 88–90% amino acid identity with other Ftf1 paralogs (Supplementary Fig. 22), yet only *FTF1a-1* and *FTF1a-3* contribute to virulence, indicating functional divergence among these highly similar paralogs encoded within AC compartments.

### FTF1a-1 regulates virulence beyond SIX effector genes

Previous studies have shown that members of the *FTF* family in 4287 regulate the expression of *SIX1*, *SIX3*, and *SIX6*^27,28^, which are located in the L15-3, L15-2, and L14-5 segments of MAFF 103036, respectively. To investigate whether *FTF1a-1* regulates *SIX* gene expression *in planta*, we quantified the expression levels of *SIX* genes in tomato roots at 7 dpi in mutants lacking *FTF1a-1*. *SIX3* expression was significantly reduced in all *FTF1a-1*-deficient mutants (ΔL16-2-1, ΔL16-2-1-1, and Δ*FTF1a-1*) compared with the wild-type, whereas reduced *SIX1* expression was observed only in the Δ*FTF1a-1* mutant (Supplementary Fig. 19). In contrast, *SIX6* expression was unchanged in all tested mutants relative to the wild-type (Supplementary Fig. 19). Strikingly, despite the full restoration of virulence observed in the complemented ΔL16-2-1-1: *FTF1a-1* strain, *SIX* expression remained lower than in the wild-type, possibly because *FTF1a-1* expression was only partially recovered (Supplementary Fig. 19). Therefore, the reduced expression of these *SIX* genes is unlikely to be the primary cause of the attenuated virulence in *FTF1a-1*-deficient mutants, suggesting that *FTF1a-1* regulates virulence-related genes other than the *SIX* effector genes.

## Discussion

The compartmentalization of fungal genomes into core and ACs represents a major evolutionary strategy driving host-specific pathogenicity; however, systematically dissecting the functional landscape of these highly repetitive and plastic regions has long remained an experimental bottleneck. We overcame this limitation by establishing a genome-editing platform that enables targeted large-scale chromosomal deletions across the AC regions in Fol. We successfully deleted 36 regions—including a segment as large as 740 kb—thereby generating a comprehensive deletion library covering approximately 99% of the targeted AC regions. This systematic mapping approach provides a post-genomic framework, shifting the current paradigm in fungal functional genomics from single-gene characterization to chromosome-scale analysis of plastic genomic regions.

Through systematic chromosome-wide screening, we successfully narrowed the genomic regions governing virulence in MAFF 103036 to seven discrete AC segments (Fig. 2d–h and Supplementary Figs. 10d–h and 11d–h). Although stepwise dissection of the L16-2 segment led to the identification of *FTF1a-1*, the remaining six AC segments still contain unresolved candidate genes responsible for virulence. For ΔL14-5, ΔL15-2, and ΔL15-3, the severe reduction in virulence is consistent with the presence of well-characterized *SIX* effector genes in these segments (Fig. 3a, f, g)^16,18,20–22^, but several uncharacterized genes within these segments showed elevated expression under *in planta* conditions (Fig. 3a, f, g and Supplementary Table 3). Similarly, in the remaining segments (L14-8, L14-10, and L14-12), which lack known virulence genes, up-or down-regulated genes during plant colonization were identified (Fig. 3b, c, e and Supplementary Table 3). The other putative AC regions which could not be tested in this study contained only 28 genes, several of which were upregulated under *in planta* conditions (Fig. 3d, h, Supplementary Fig. 23a–h, and Supplementary Tables 3 and 4). Although most of these genes encode proteins of unknown function, our systematic dissection substantially narrows the set of candidate virulence genes in the ACs, providing a defined starting point for their functional characterization in the future.

The identification of *FTF1a-1* as a dominant virulence regulator represents the central mechanistic insight of this study. Previous studies in *F. oxysporum* f. sp. *phaseoli* reported that isolates carrying a higher *FTF1* copy number exhibit higher overall *FTF1* transcript abundance and enhanced virulence, although the contributions of individual paralogs remained unresolved^27,33,34^. By systematically deleting each paralog, we resolved these individual contributions in Fol and found that the virulence function is largely attributable to a single paralog, *FTF1a-1*, rather than to the *FTF1* family as a whole. Although the promoters of *FTF1a-2* and *FTF1a-3* are 90% and 87% identical to that of *FTF1a-1* over the upstream 340 bp, respectively (Supplementary Fig. 24)^27^, *FTF1a-2* was not expressed during host colonization (Supplementary Table 1). By contrast, the remaining *FTF1a* and *FTF1c* paralogs showed similar expression patterns under *in planta* conditions in our RNA-seq data (Supplementary Table 1), as also reported in *F. oxysporum* f. sp. *phaseoli*^34^. Despite these similar expression patterns, the paralogs differed in their contribution to virulence; only the loss of *FTF1a-1* caused a severe reduction in virulence, even in the presence of the other paralogs (Supplementary Fig. 21a–i). Together, our results indicate that *FTF1a-1* is functionally distinct from the other paralogs and plays a dominant role in virulence.

The functional non-redundancy of *FTF1a-1* is supported by domain-level amino acid sequence divergence. Although Ftf1 paralogs of MAFF 103036 share 88–90% overall amino acid identity, sequence alignments revealed divergence within two functional domains: the zinc-finger-like DNA-binding region (approximately residues 175–235) and the C-terminal regulatory activation domains (approximately after residue 800) that interact with the transcriptional machinery^28,35^ (Supplementary Fig. 22). The distinct contributions of *FTF1a-1* and *FTF1a-3* to Fol virulence are likely attributable to functional divergence in these domains. Similar analogous diversification has also been described in *Candida albicans*, where duplicated transcription factors acquire distinct functions through altered DNA-binding specificities and cofactor associations^36,37^. Previous transcriptional profiling of *FTF1a-1*-overexpressing 4287 showed that *FTF1a-1* upregulates the core chromosome-encoded transcription factor *FOXG_04965*, which is required for pathogenicity, together with numerous AC-encoded genes^28^. These include uncharacterized small secreted protein genes linked to (partial) miniature impala transposable elements, and secondary metabolite biosynthetic gene clusters involved in C-compound and carbohydrate metabolism^28^. Several genes homologous to these are located within the virulence-associated AC regions identified in this study (Supplementary Table 5), suggesting that they may represent downstream targets of *FTF1a-1* in MAFF 103036.

Previous studies have highlighted the multifaceted roles of fungal ACs beyond pathogenicity. For instance, adaptive mutations in an AC-encoded gene in 4287 impacted not only virulence but also colony growth, conidiation, and quorum sensing^38^. Similarly, disruption of an AC-encoded gene in *F. oxysporum* f. sp. *cepae* led to increased colony growth and conidiation^39^, while the ACs of *F. oxysporum* f. sp. *cubense* contribute to tolerance against hyperosmotic and cell wall stresses^40^. Together, these observations suggest that AC regions confer broader adaptive advantages, both during host infection and throughout diverse physiological processes outside the host. Consequently, our chromosomal deletion library also provides a valuable resource for exploring how fungi leverage plastic genomic compartments to survive diverse environmental and evolutionary constraints.

## Methods

### Fungal isolate and culture conditions

Fol race 1 isolate MAFF 103036 was used throughout this study. The isolate was cultured on potato dextrose agar (PDA; Nissui, Japan) or YG agar medium [0.5% (w/v) yeast extract, 2% (w/v) glucose, 1.5% (w/v) agar] at 28 °C in the dark for 3–5 days. Long-term storage of the isolate was performed in 25% (w/v) glycerol at −80 °C. For microconidia (bud-cell) production, mycelial plugs derived from PDA cultures were inoculated into potato dextrose broth [PDB; potato extract containing 0.5% (w/v) dextrose] and incubated at 26 °C with shaking at 120 revolutions per minute (rpm) for 3–5 days. To determine colony growth rates, a mycelial plug was punched out using a cork borer and placed onto a PDA plate, and colony diameters were measured after 4 days of incubation.

### Construction of donor vectors and CRISPR/Cas9 vectors

Donor vectors for large-scale deletion of AC segments and for single-gene disruption were constructed by amplifying homology arms of the target segments or genes via PCR. The amplified upstream homologous arm was cloned into pMK-dGFP–which harbors the *Hph* cassette^41^– utilizing either the *Kpn*I, *Xho*I, *Sph*I or *Afe*I restriction sites to generate pMK-FL, or alternatively, the amplified downstream homology arm was inserted between the *Hind*III, *EcoR*V, *Spe*I, *Not*I, or *Sac*I sites to generate pMK-FR. Subsequently, the remaining homology arm was inserted into pMK-FL or pMK-FR using the corresponding restriction sites to generate the final donor vector, pMK-FL-FR.

For construction of the donor vector used to generate the ΔL16-2-1-1: *FTF1a-1* complementation strain, primers were designed to amplify the *FTF1a-1* coding region together with the predicted 1,500-bp upstream promoter region and 1,500-bp downstream terminator region. The amplified fragments were inserted between the *EcoR*I and *EcoR*V sites of the pII99 vector containing the *NPTII* (geneticin resistance gene) cassette.

To ensure efficient and specific induction of DNA double-strand breaks by CRISPR/Cas9, all potential gRNA target sequences were designed within open reading frames and screened by local BLAST against the Fol genome to minimize potential off-target sites. Predicted off-target sites, if identified, were PCR-amplified and verified by Sanger sequencing. CRISPR/Cas9 expression vectors were constructed by annealing the selected oligonucleotide primer sets according to previously described procedures^42,43^. The annealed oligonucleotides were inserted into the two *Esp3*I sites of the improved pCRISPR/Cas9 vector for *F. oxysporum*^40^ via Golden Gate cloning following previously described procedures^42,43^. All primers used for donor vector and CRISPR/Cas9 vector construction are listed in Supplementary Table 2.

### Protoplast preparation and transformation of Fol

Protoplast preparation and transformation of Fol were performed according to a previously described method^25^, with minor modifications. Briefly, small mycelial plugs derived from PDA cultures were transferred to 25 mL of YG liquid medium and incubated at 28 °C for 1 day with shaking at 120 rpm. One milliliter of this culture was transferred into 100 mL of fresh YG liquid medium and incubated under the same conditions for an additional day. The culture was divided into two aliquots, and mycelia were collected by centrifugation at 5800 *× g* for 10 min. The mycelial pellets were washed once with 20 mL of sterile water and twice with 10 mL of 1.2 M MgSO4 with centrifugation at 5800 *× g* for 10 min after each wash. Mycelia were then treated with a cell-wall-degrading enzyme mixture containing 20 mg/mL Yatalase (Takara Bio, Japan), 20 mg/mL Lysing enzyme (Sigma, USA), and 25 mg/mL Extralyze (LAFFORT, France) in 1.2 M MgSO_4_ for 5 h at 28 °C. Protoplasts were washed twice with STC buffer [1.2 M sorbitol, 50 mM CaCl_2_, 10 mM Tris-HCl pH 7.5] and adjusted to 5 × 10⁶ cells/mL.

For gene disruptions, 4 µg of a CRISPR/Cas9 vector and 3 µg of a pMK-dGFP-based donor vector were mixed (final volume ≤ 5 µL). For large-scale deletions, 2 µg each of two CRISPR/Cas9 vectors and 3 µg of a pMK-dGFP-based donor vector were mixed (final volume ≤ 5 µL). For genetic complementation of the ΔL16-2-1-1 mutant with *FTF1a-1*, 4 µg of a pII99-based donor vector was mixed with the protoplasts (final volume ≤ 5 µL). Subsequently, 50 µL of the protoplast suspension was added to each mixture, followed by a gentle overlay of 50 µL of 60% (w/v) PEG 4000 (Wako, Japan). The mixture was incubated at room temperature for 20 min. After incubation, 1.0 mL of STC buffer was added, and the mixture was centrifuged at 3800 *× g* for 5 min at room temperature. The supernatant was discarded, and the pellet was resuspended in 250 µL of STC buffer. The suspension was poured into a sterile petri dish, mixed with 10 mL of molten RM medium [0.5% [w/v] yeast extract, 34.2% [w/v] sucrose, and 0.8% agar], gently mixed, and allowed to solidify at room temperature. Finally, YG medium supplemented with hygromycin B (100 µg/mL) or PDA medium supplemented with geneticin (150 µg/mL) was overlaid onto the RM agar for the selection of transformants. All transformants were verified by PCR, and the primers used for PCR are listed in Supplementary Table 2.

### Plant growth conditions and tomato inoculation tests

Tomato seeds (*S. lycopersicum* cv. Ponderosa; Noguchi Seed, Japan) were planted at a density of 5–6 seeds per pot in autoclaved soil (Kumiai Nippi Engeibaido No. 1; Nihon Hiryo, Japan) using plastic pots and grown in a greenhouse at 24 °C. Fourteen-day-old tomato seedlings were utilized for subsequent experiments. For inoculation, Fol and its transformants were cultured in PDB. Bud-cells were collected by centrifugation at 4000 *× g* for 5 min, resuspended in sterile water to a final concentration of 1 × 10^8^ cells/mL, and used as an inoculum. Disease assays were performed via soil-drench inoculation. For each plastic pot, two seedlings were retained. Each seedling was inoculated by applying 2 mL of bud-cells suspension to the base of the stem. Disease symptoms were evaluated at 30–42 days post-inoculation (dpi) based on both external and internal symptoms. External symptoms were scored according to the degree of leaf yellowing, wilting, or mortality using the following disease index: 0, yellowing or wilting of fewer than two true leaves; 1, yellowing or wilting of three true leaves; 2, yellowing or wilting of four or more true leaves; 3, yellowing or wilting of the upper leaves; 4, plant death. Internal disease symptoms were evaluated by sectioning the stem near the root interface and scoring the percentage of vascular bundle browning as follows: 0, no symptoms; 1, 25% browning; 2, 50% browning; 3, 75% browning; 4, 100% browning.

For RNA extraction, plant samples were prepared using a root-dip inoculation. Soil was removed from the roots by washing with sterile water, and the roots were then immersed in the bud-cells suspension (1 × 10^7^ cells/mL) for a minimum of 5 min. Root tissues were collected at 7 dpi, washed extensively with sterile water, and stored at −80 °C until RNA extraction.

### DNA extraction, whole-genome sequencing, assembly, and BUSCO analysis

Genomic DNA was extracted from bud-cells using hexadecyltrimethylammonium bromide and purified with Genomic-tip 100/G kit (QIAGEN, Germany) as described in the 1000 Fungal Genomes Project. Whole-genome sequencing was performed utilizing a PacBio Sequel SMRT cell (PacBio, USA). De novo genome assembly was conducted using the Hierarchical Genome Assembly Process (HGAP) v4 within SMRT Link v10.2.0.133434, following a previously described procedure^44^. The expected genome size parameter was configured to 60 Mb, with all other parameters maintained at their default settings. The completeness of the resulting genome assembly was assessed using BUSCO v6.0.0^45^ against the Sordariomycetes_odb12 dataset of OrthoDB.

### RNA-seq, mapping, and quantification

Total RNA was extracted using the RNeasy Plant Mini Kit (QIAGEN, Germany) according to the manufacturer’s instructions. Library preparation and sequencing (150-bp paired-end reads) were performed by Rhelixa (Tokyo, Japan). Raw sequence reads were quality-filtered using fastp v1.0.1^46^; the high-quality reads were subsequently aligned to the reference genome assembly using HISAT2^47^. Quantifications of the mapped reads for each predicted gene locus were performed using featureCounts^48^.

### Repeat masking, gene prediction, and genome alignment

Repetitive sequences within the assembled contigs were soft-masked using RepeatMasker v4.2.3, utilizing a custom repeat library generated by combining datasets from Dfam 3.9^49^ and Repbase^50^. The soft-masked genomic sequences, along with both *in vitro* and *in planta* RNA-seq reads, were subjected to *ab initio* and evidence-based gene prediction using the BRAKER v3.0.8^51–55^ pipeline incorporating DIAMOND v0.9.24^56^, GeneMark-ETP^57^, GffRead^58^, Spaln^59,60^, StringTie2^61^, and TSEBRA^52^. Whole-genome alignments were performed using the “nucmer” module in MUMmer v4.0.1^62^ with the options --maxmatch and -l 500, with all other parameters maintained at their default settings. Aligned sequences were visualized using Circos v0.69-10^63^.

### Functional annotation

Functional annotation was performed utilizing Blast2GO v6.0.3^64^. Predicted protein sequences were queried against the NCBI non-redundant (nr) protein database via BLASTP with an E-value cut-off of 1 × 10^-3^, and scanned for conserved protein signatures using InterProScan, both as implemented within the Blast2GO platform. Gene Ontology terms and functional descriptions were then assigned by combining the BLASTP and InterProScan results. Gene product names were manually curated where necessary.

### RNA extraction, cDNA synthesis, and gene expression by RT–qPCR analysis

Total RNA was extracted using the ISOSPIN Plant RNA kit (NIPPON Gene, Japan) according to the manufacturer’s instructions, including on-column DNase I treatment to remove residual genomic DNA. First-strand cDNA was synthesized from 400 ng of total RNA using ReverTra Ace (Toyobo, Japan). The resulting cDNA was diluted 10-fold, and a 2 µL aliquot was utilized as a template in a 25 µL reaction mixture containing GoTaq qPCR Master Mix (Promega, USA). qPCR was performed using a Thermal Cycler Dice Real Time System III (Takara Bio, Japan) under the following conditions: 95 °C for 2 min, followed by 40 cycles of 95 °C for 5 s and 60 °C for 30 s. Relative gene expression levels were calculated using the ΔΔ*Ct* method^65^. Previously reported primers were used for *EF1a, SIX1, SIX3, SIX6,* and *FTF1a-1*^28,32,66^. All primers used for qPCR are listed in Supplementary Table 2.

### Statistics and reproducibility

Statistical analyses were performed using R (version 4.5.2). Disease index data were analyzed using two-sided Mann–Whitney *U*-tests or Kruskal–Wallis tests followed by Dunn’s multiple-comparison tests, as indicated in the corresponding figure legends. Where multiple comparisons against the wild type were performed within a single experiment, *P* values were adjusted using the Benjamini–Hochberg method. Colony growth assay data and RT–qPCR data were analyzed by one-way analysis of variance (ANOVA) followed by Tukey’s multiple-comparison test. The numbers of biologically independent replicates and the statistical tests used are specified in the corresponding figure legends.

## Data availability

The whole-genome and RNA-sequencing data generated in this study have been deposited in the DNA Data Bank of Japan (DDBJ) database under BioProject accession number PRJDB42575. The genome assembly and RNA-seq reads are available under BioSample accession numbers SAMD01933835, SAMD01936993, and SAMD01936994.

## ACKNOWLEDGMENTS

This work was supported in part by the Japan Society for the Promotion of Science (JSPS) KAKENHI Grant-in-Aid for Scientific Research (A) (Grant numbers 19H00939 and 23H00331) for TS.A., S.A. and TA.A., and Grant-in-Aid for Challenging Exploratory Research (Grant number 18K19218) for TA.A.; Japan Science and Technology Agency (JST) PRESTO (Grant number JPMJPR16O1) and FOREST (Grant number JPMJFR2101) for S.A.; the Science and Technology Research Partnership for Sustainable Development (SATREPS; grant number 22571054) funded jointly by the Japan Science and Technology Agency (JST) and Japan International Cooperation Agency (JICA) for TS.A.; and the FLOuRISH Fellowship Program of TUAT to M.Y. We used ChatGPT (OpenAI, GPT-5.5) and Claude (Anthropic, Claude Sonnet 4) for the grammatical refinement of the manuscript with conversation history for model improvement disabled.

## Author Contributions

TA.A. and TS.A. conceived and designed the study. M.Y. performed most of the experiments. TA.A., TS.A., T.K., K.K., and M.Y. analyzed the data. TA.A., TS.A., and M.Y. drafted the manuscript. S.A. and H.S. performed the whole-genome sequencing and RNA-seq analyses, and S.A. provided technical expertise. TA.A., K.K., and TS.A. supervised the study. All authors discussed the results, reviewed the manuscript, and approved the final version.

