## Supplemental Figures for "Comprehensive functional mapping of accessory chromosomes identifies a dominant virulence-regulator paralog in tomato wilt pathogen": Supplementary information.pdf

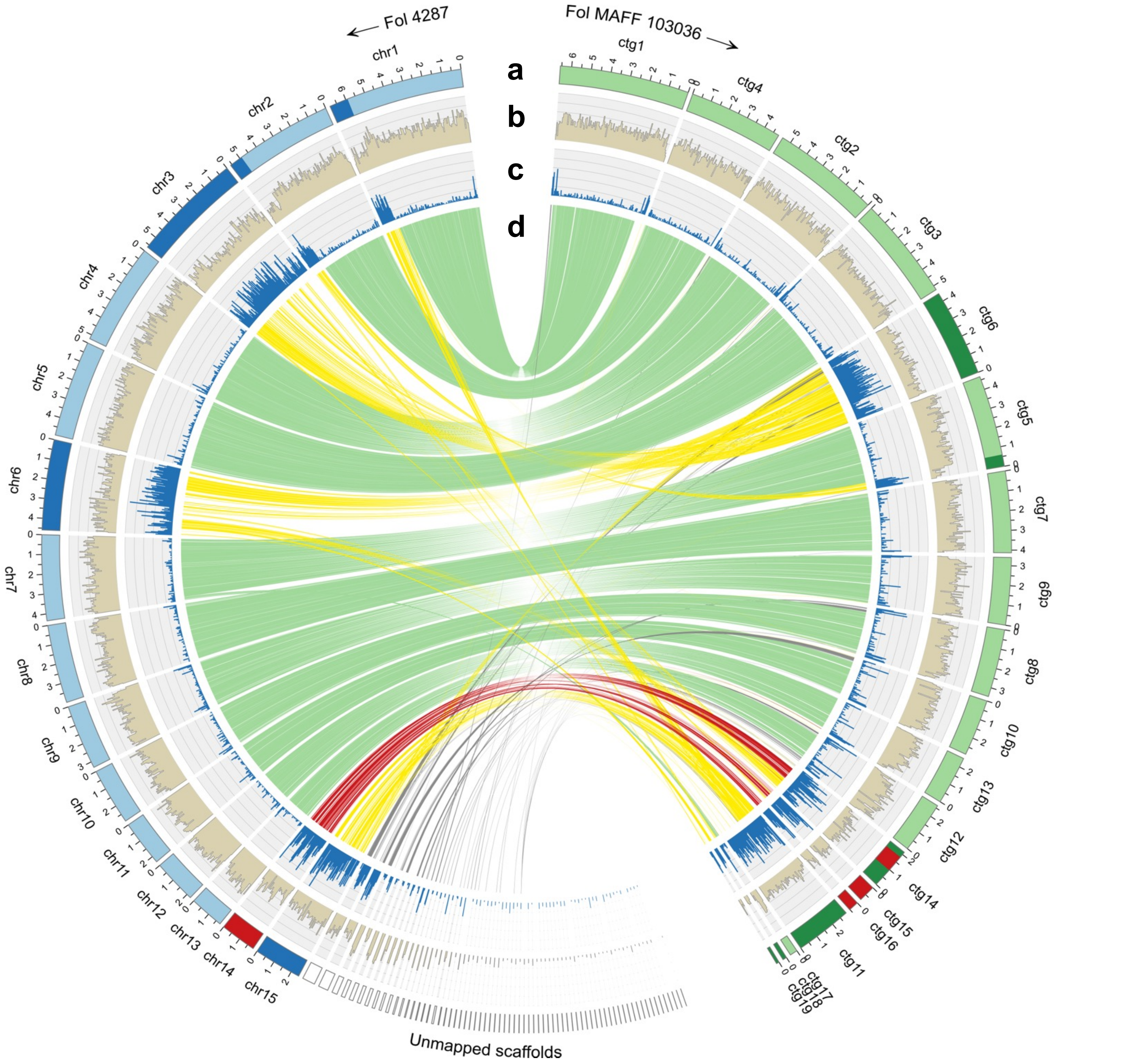


**Supplementary Figure 1: Comparison of the whole-genome assemblies of 4287 and MAFF 103036. a** Chromosomes of 4287 and contigs of MAFF 103036. In 4287, chromosomal regions were annotated based on Ma et al. (2010): core chromosomal regions, AC regions, and the pathogenicity chromosome (chromosome 14) are shown in light blue, dark blue, and red, respectively. Uncolored boxes indicate unmapped scaffolds. In MAFF 103036, contigs corresponding to core chromosomal regions, AC regions, and chromosome 14 are shown in light green, dark green, and red, respectively. Abbreviations: chr, chromosome; ctg, contig. **b** Gene density in 50-kb windows. **c** Repeat density in 50-kb windows. **d** Alignments between 4287 and MAFF 103036, as determined by nucmer. Alignment lines corresponding to chromosome 14 of 4287 are shown in red, those corresponding to the ACs other than chromosome 14, and those corresponding to the core chromosome are shown in light green.

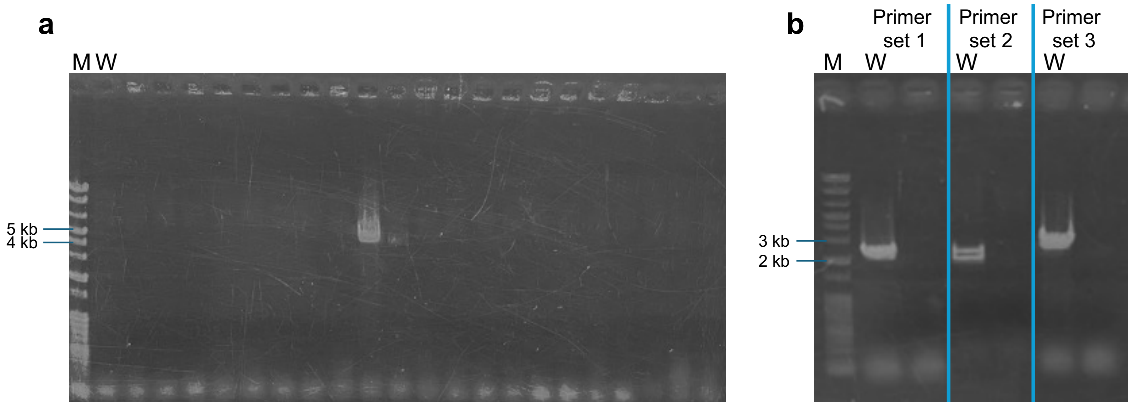


**Supplementary Figure 2: PCR-based validation of the large-scale deletion of the L15-3 segment.** Representative agarose gel images confirming the targeted deletion of the L15-3 segment. **a** A 4,660-bp PCR product was detected only in the deletion mutant. **b** Verification of deletion of the internal region using three primer sets located within the L15-3 segment in the transformant shown in (**a**). PCR products of 2,247, 2,009, and 2,722 bp were expected from primer sets 1, 2, and 3, respectively, in the presence of the intact wild-type locus, whereas no amplification was observed in the deletion mutants. M, molecular weight marker; W, wild-type.

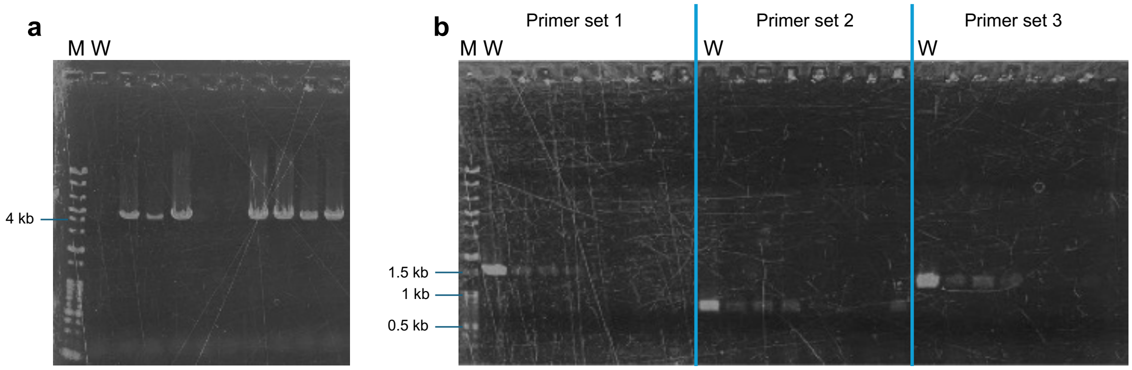


**Supplementary Figure 3: PCR-based validation of the large-scale deletion of the L5-1 segment.** Representative agarose gel images confirming the targeted deletion of the L5-1 segment. **a** A 4,052-bp PCR product was detected only in the targeted deletion mutants. **b** Verification of deletion of the internal region using three primer sets located within the L5-1 segment in the transformants shown in (a). PCR products of 1,501 bp, 776 bp, and 1,244 bp were expected from primer sets 1, 2, and 3, respectively, in the presence of the intact wild-type locus, whereas no amplification was observed in the deletion mutants. M, molecular weight marker; W, wild-type.

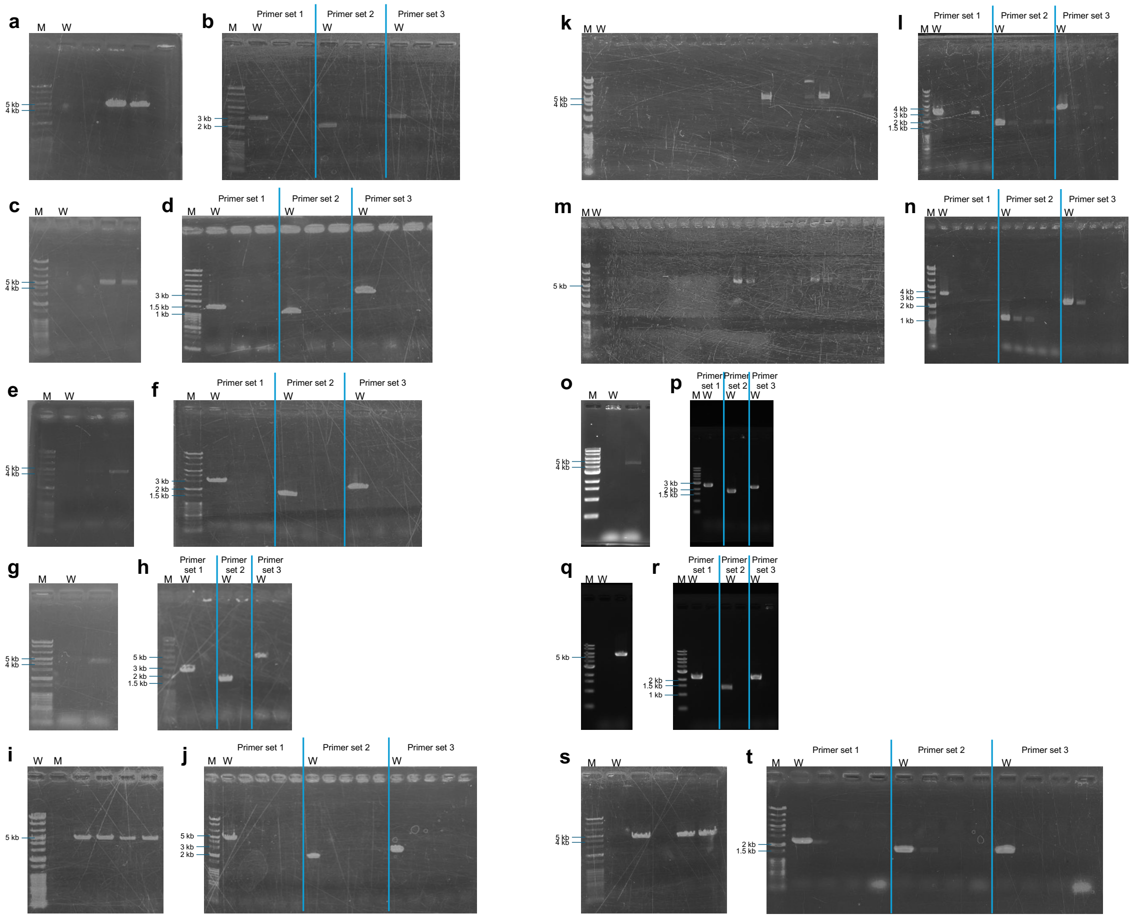


**Supplementary Figure 4: PCR-based validation of the large-scale deletion of the L6-1 to L6-10 segments.** Representative agarose gel images confirming the targeted deletion of the L6-1 to L6-10 segments. **a,c,e,g,i,k,m,o,q,s** PCR products were detected only in the targeted deletion mutants, yielding fragments of 4,663 bp (L6-1; a), 4,511 bp (L6-2; c), 4,502 bp (L6-3; e), 4,815 bp (L6-4; g), 5,116 bp (L6-5; i), 4,609 bp (L6-6; k), 5,706 bp (L6-7; m), 4,744 bp (L6-8; o), 5,508 bp (L6-9; q), and 4,502 bp (L6-10; s). **b,d,f,h,j,l,n,p,r,t** Verification deletion of the internal region using three primer sets located within each targeted segment in the transformants shown in (a, c, e, g, i, k, m, o, q and s). PCR products of 3,070, 1,991, and 2,895 bp (L6-1; b); 1,447, 1,092, and 2,927 bp (L6-2; d); 2,927, 1,776, and 2,645 bp (L6-3; f); 2,645, 1,569, and 5,157 bp (L6-4; h); 5,157, 2,114, and 2,907 bp (L6-5; j); 2,907, 1,644, and 3,765 bp (L6-6; l); 3,765, 1,127, and 2,521 bp (L6-7; n); 2,521, 1,697, and 2,264 bp (L6-8; p); 2,264, 1,420, and 2,294 bp (L6-9; r); and 2,294, 1,628, and 1,676 bp (L6-10; t) were expected from primer sets 1, 2, and 3, respectively, in the presence of the intact wild-type locus, whereas no amplification was observed in the deletion mutants. M, molecular weight marker; W, wild-type.

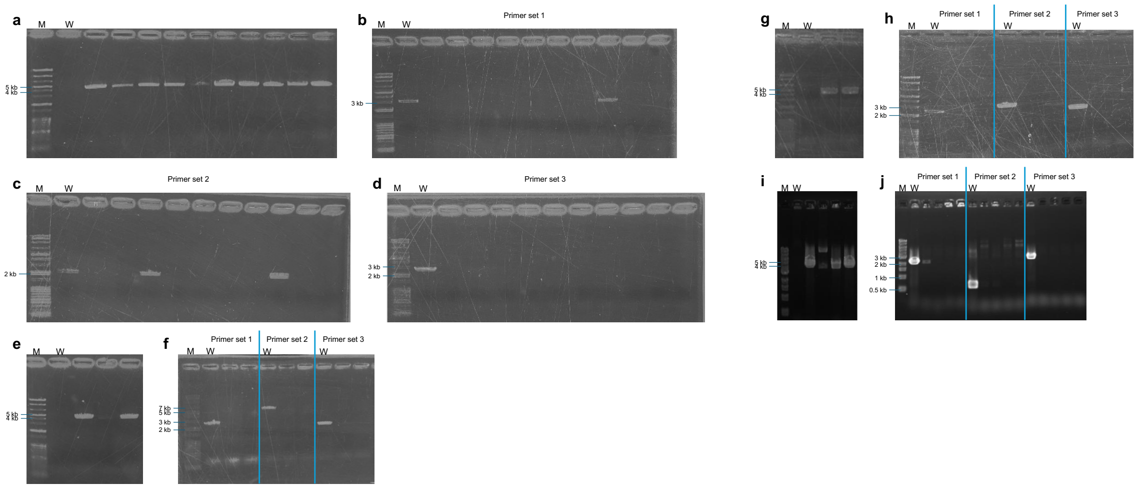


**Supplementary Figure 5: PCR-based validation of the large-scale deletion of the L11-1 to L11-4 segments.** Representative agarose gel images confirming the targeted deletion of the L11-1 to L11-4 segments. **a,e,g,i** PCR products were detected only in the targeted deletion mutants, yielding fragments of 4,514 bp (L11-1; a), 4,502 bp (L11-2; e), 4,484 bp (L11-3; g), and 4,634 bp (L11-4; i). **b–d,f,h,j** Verification deletion of the internal regions using three primer sets located within each targeted segment in the transformants shown in (a, e, g and i). PCR products of 3,093, 2,248, and 2,663 bp (L11-1; b–d); 2,663, 6,581, and 2,463 bp (L11-2; f); 2,524, 2,882, and 2,244 bp (L11-3; h); and 2,244, 618, and 2,605 bp (L11-4; j) were expected from primer sets 1, 2, and 3, respectively, in the presence of the intact wild-type locus, whereas no amplification was observed in the deletion mutants. M, molecular weight marker; W, wild-type.

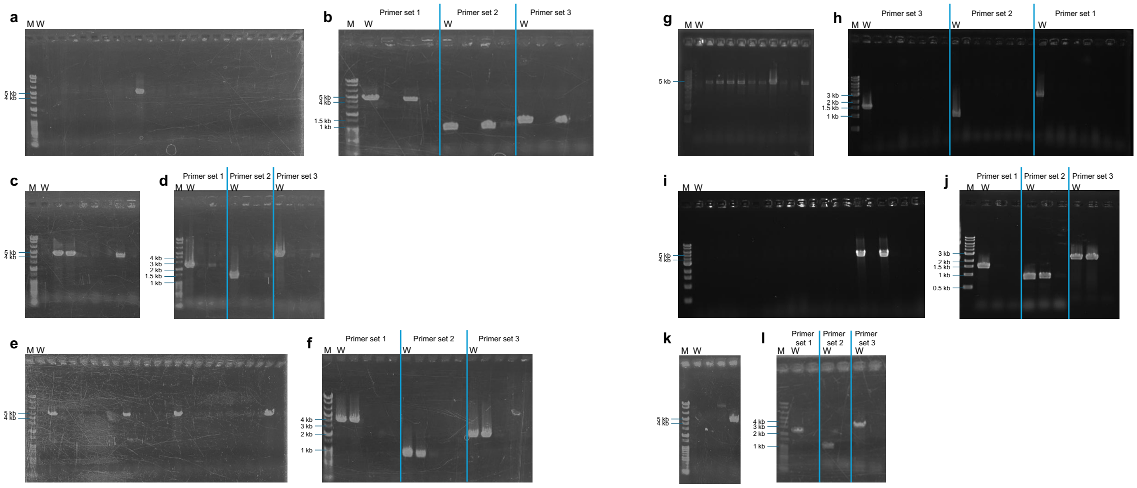


**Supplementary Figure 6: PCR-based validation of the large-scale deletion of the L14-1 to L14-6 segments.** Representative agarose gel images confirming the targeted deletion of the L14-1 to L14-6 segments. **a,c,e,g,i,k** PCR products were detected only in the targeted deletion mutants, yielding fragments of 4,573 bp (L14-1; a), 4,518 bp (L14-2; c), 4,780 bp (L14-3; e), 4,996 bp (L14-4; g), 4,535 bp (L14-5; i), and 4,671 bp (L14-6; k). **b,d,f,h,j,l** Verification deletion of the internal region using three primer sets located within each targeted segment in the transformants shown in (a, c, e, g, I and k). PCR products of 4,170, 1,046, and 1,312 bp (L14-1; b); 2,326, 1,304, and 3,965 bp (L14-2; d); 3,965, 1,014, and 2,521 bp (L14-3; f); 2,521, 826, and 1,551 bp (L14-4; h); 1,449, 868, and 2,368 bp (L14-5; j); and 2,368, 1,000, and 3,117 bp (L14-6; l) were expected from primer sets 1, 2, and 3, respectively, in the presence of the intact wild-type locus, whereas no amplification was observed in the deletion mutants. M, molecular weight marker; W, wild-type.

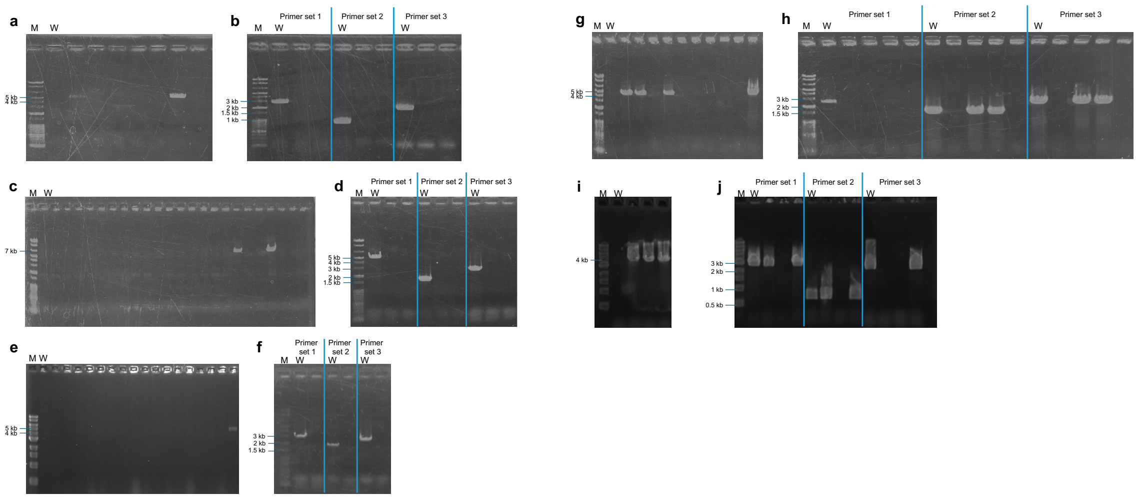


**Supplementary Figure 7: PCR-based validation of the large-scale deletion of the L14-7 to L14-12 segments, excluding L14-11.** Representative agarose gel images confirming the targeted deletion of the L14-7 to L14-12 segments, excluding L14-11. **a,c,e,g,i** PCR products were detected only in the targeted deletion mutants, yielding fragments of 4,513 bp (L14-7; a), 7,197 bp (L14-8; c), 4,759 bp (L14-9; e), 4,674 bp (L14-10; g), and 4,064 bp (L14-12; i). **b,d,f,h,j** Verification deletion of the internal region using three primer sets located within each targeted segment in the transformants shown in (a, c, e, g and i). PCR products of 2,795, 885, and 1,871 bp (L14-7; b); 5,044, 1,736, and 2,846 bp (L14-8; d); 2,846, 1,861, and 2,566 bp (L14-9; f); 2,566, 1,531, and 2,478 bp (L14-10; h); and 2,977, 708, and 2,639 bp (L14-12; j) were expected from primer sets 1, 2, and 3, respectively, in the presence of the intact wild-type locus, whereas no amplification was observed in the deletion mutants. M, molecular weight marker; W, wild-type.

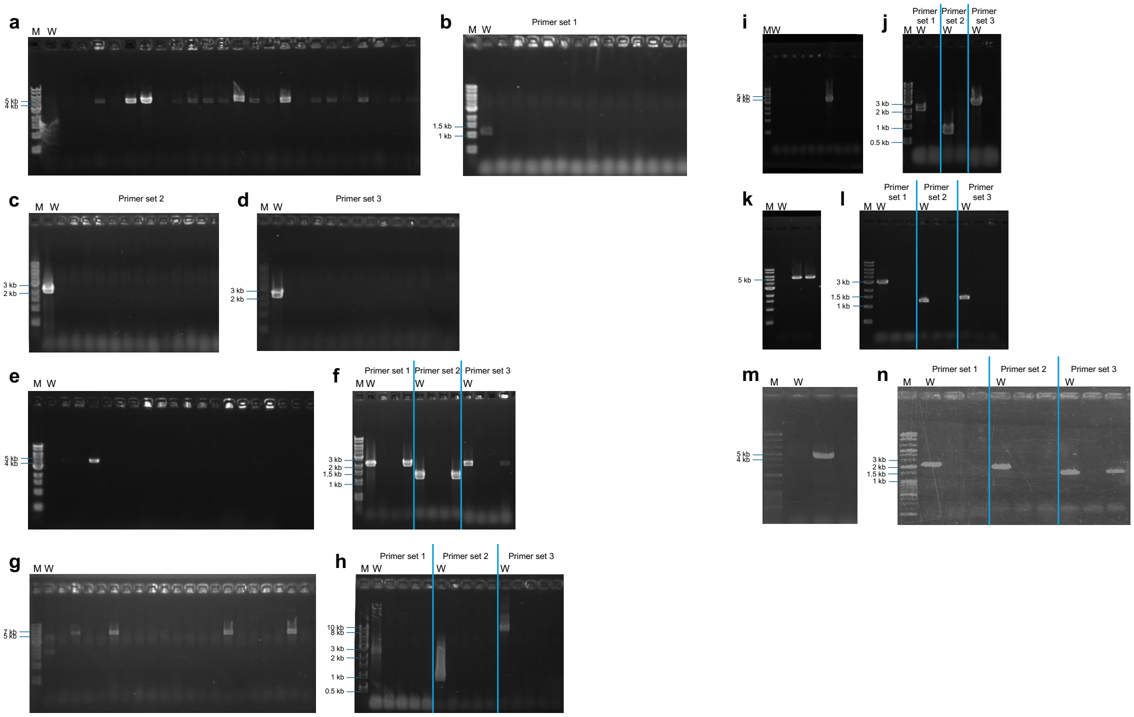

**Supplementary Figure 8: PCR-based validation of the large-scale deletion of the L15-1 to L15-8 segments, excluding L15-3 and L15-5.** Representative agarose gel images confirming the targeted deletion of the L15-1 to L15-8 segments, excluding L15-3 and L15-5. **a,e,g,i,k,m** PCR products were detected only in the targeted deletion mutants, yielding fragments of 4,559 bp (L15-1; a), 4,662 bp (L15-2; e), 6,542 bp (L15-4; g), 4,450 bp (L15-6; i), 5,228 bp (L15-7; k), and 4,407 bp (L15-8; m). **b–d,f,h,j,l,n** Verification deletion of the internal region using three primer sets located within each targeted segment in the transformants shown in (a, e, g, i, k and m). PCR products of 1,153, 2,323, and 2,224 bp (L15-1; b–d); 2,224, 1,256, and 2,247 bp (L15-2; f); 2,722, 878, and 9,168 bp (L15-4; h); 2,243, 807, and 2,974 bp (L15-6; j); 2,974, 1,277, and 1,445 bp (L15-7; l); and 2,222, 2,080, and 1,504 bp (L15-8; n) were expected from primer sets 1, 2, and 3, respectively, in the presence of the intact wild-type locus, whereas no amplification was observed in the deletion mutants. M, molecular weight marker; W, wild-type.

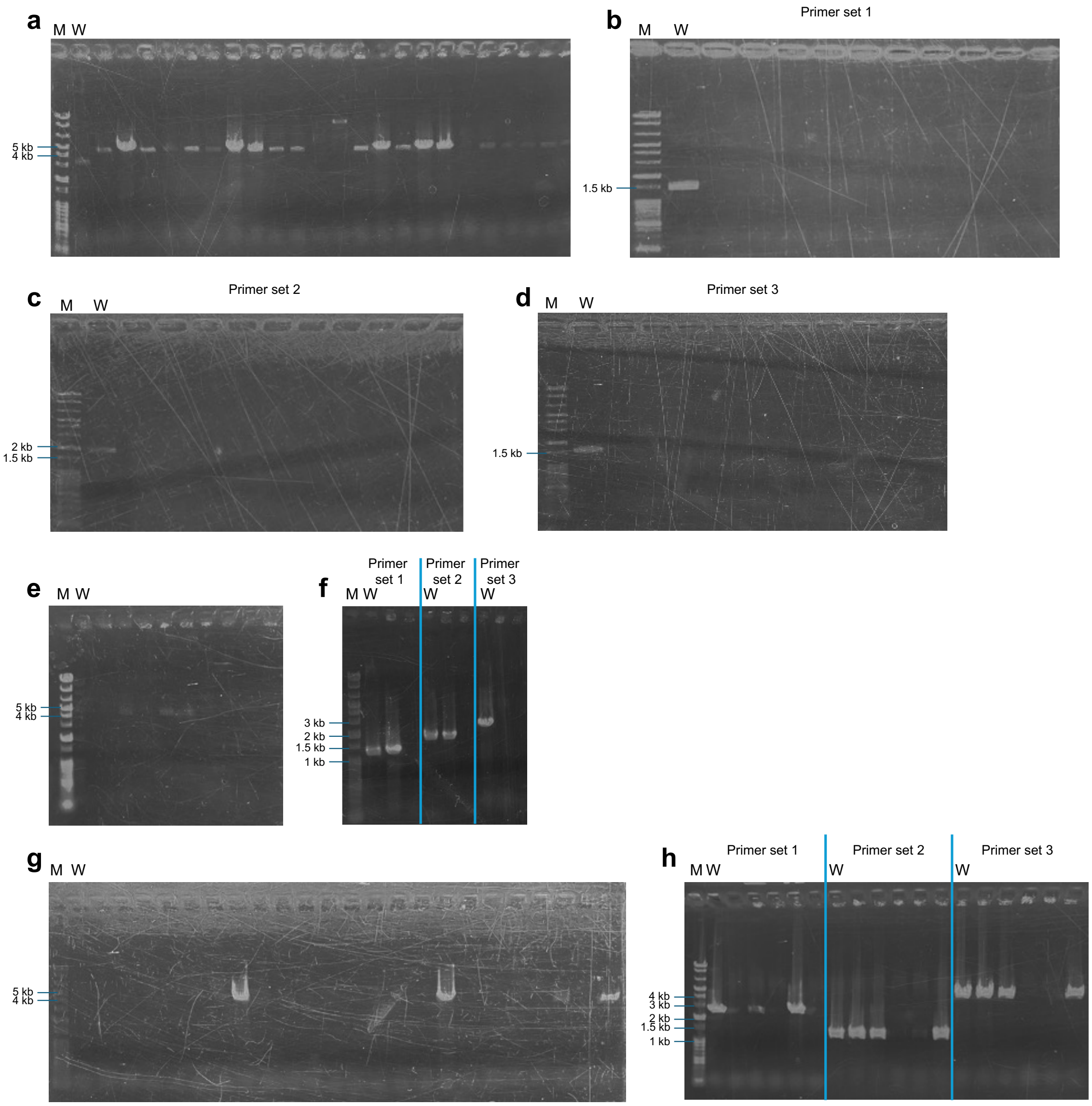


**Supplementary Figure 9: PCR-based validation of the large-scale deletion of the L16-1 to L16-3 segments.** Representative agarose gel images confirming the targeted deletion of the L16-1 to L16-3 segments. **a,e,g** PCR products were detected only in the targeted deletion mutants, yielding fragments of 4,591 bp (L16-1; a), 4,491 bp (L16-2; e), and 4,598 bp (L16-3; g). **b–d,f,h** Verification deletion of the internal region using three primer sets located within each targeted segment in the transformants shown in (a, e and g). PCR products of 1,456, 1,830, and 1,565 bp (L16-1; b–d); 1,259, 1,779, and 2,464 bp (L16-2; f); and 2,464, 1,215, and 4,140 bp (L16-3; h) were expected from primer sets 1, 2, and 3, respectively, in the presence of the intact wild-type locus, whereas no amplification was observed in the deletion mutants. M, molecular weight marker; W, wild-type.

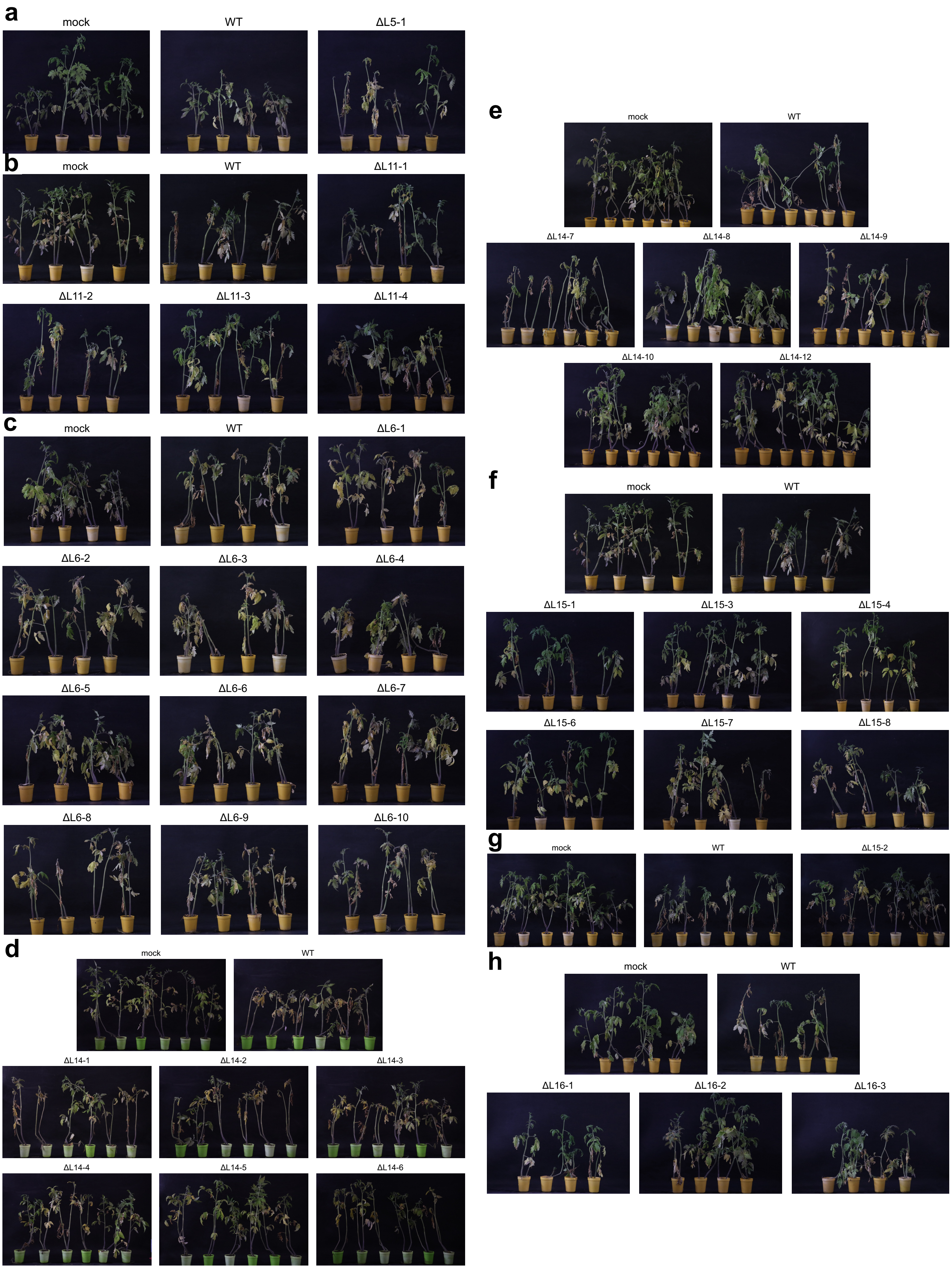
a
mock
WT
ΔL5-1
b
mock
WT
ΔL11-1
ΔL11-2
ΔL11-3
ΔL11-4
c
mock
WT
ΔL6-1
ΔL6-2
ΔL6-3
ΔL6-4
ΔL6-5
ΔL6-6
ΔL6-7
ΔL6-8
ΔL6-9
ΔL6-10
d
mock
WT
ΔL14-1
ΔL14-2
ΔL14-3
ΔL14-4
ΔL14-5
ΔL14-6
e
mock
WT
ΔL14-7
ΔL14-8
ΔL14-9
ΔL14-10
ΔL14-12
f
mock
WT
ΔL15-1
ΔL15-3
ΔL15-4
ΔL15-6
ΔL15-7
ΔL15-8
g
mock
WT
ΔL15-2
h
mock
WT
ΔL16-1
ΔL16-2
ΔL16-3

**Supplementary Figure 10: Representative external disease symptoms caused by AC deletion mutants in MAFF 103036. a–h** Representative images of tomato plants inoculated with deletion mutants corresponding to segments on contig 5 (a), contig 11 (b), contig 6 (c), the long arm of contig 14 (d), the short arm of contig 14 (e), contig 15 (f and g), and contig 16 (h). For each mutant, disease symptoms were evaluated by comparison with mock- and wild-type-inoculated plants.

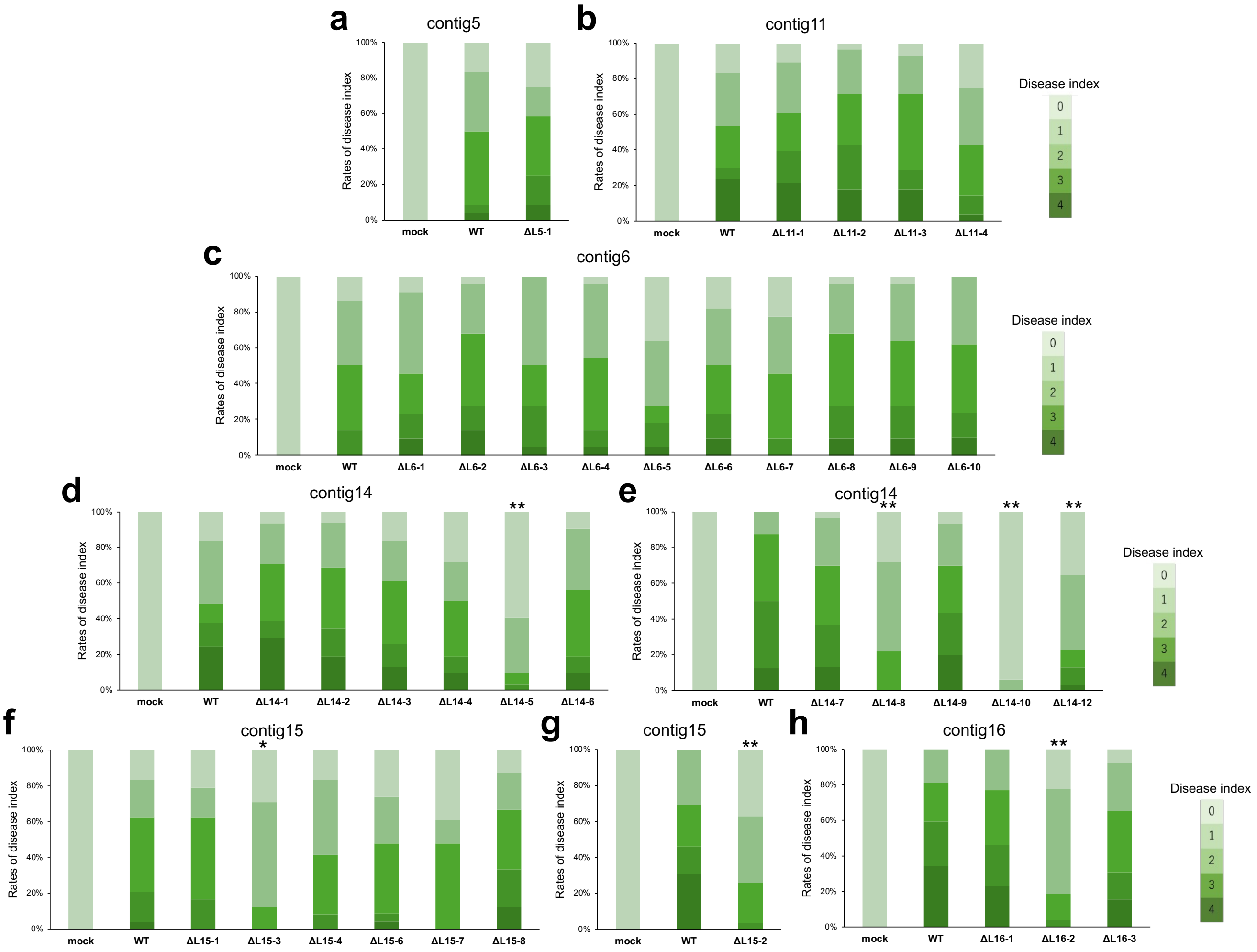


**Supplementary Figure 11: Virulence analysis of the AC deletion library in MAFF 103036 based on internal disease symptoms. a–h** Deletion mutants corresponding to segments in contig 5 (a), contig 11 (b), contig 6 (c), the long arm of contig 14 (d), the short arm of contig 14 (e), contig 15 (f and g), and contig 16 (h) were evaluated based on internal disease symptoms. Disease index was scored as described in the Methods. Asterisks indicate significant differences compared with the wild-type. (n = 21–37 biological replicates; *adjusted *P* < 0.05, **adjusted *P* < 0.01; Mann–Whitney *U*-test).

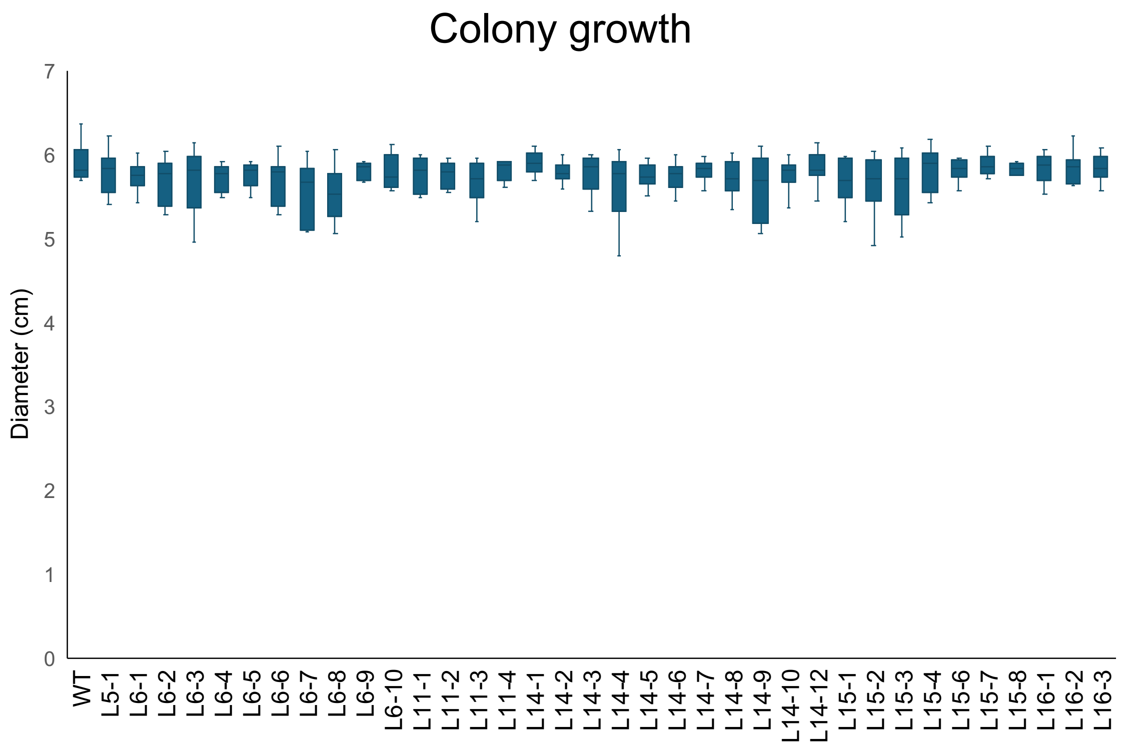


**Supplementary Figure 12: Comparison of colony growth on PDA between the wild-type and all mutants of the AC library.** Data are presented as means ± standard deviations (SD) from three biologically independent experiments, with three technical replicates each (total n = 9). Statistical significance was assessed by one-way analysis of variance (ANOVA) followed by Tukey's multiple-comparison test. No statistically significant differences were detected among the strains (adjusted $P \geq 0.05$).

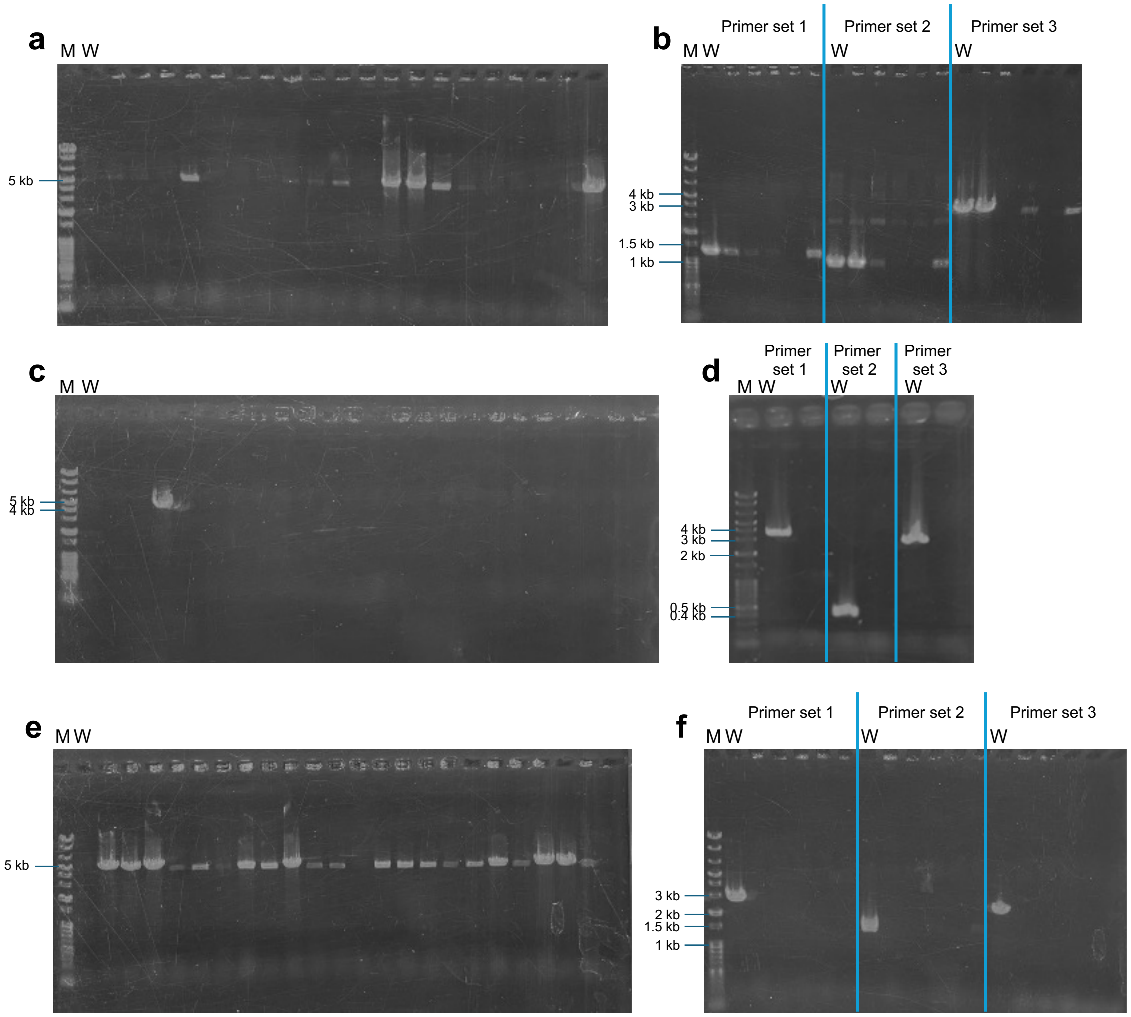


**Supplementary Figure 13: PCR-based validation of the large-scale deletion of the L16-2-1 to L16-2-3 segments.** Representative agarose gel images confirming the targeted deletion of the L16-2-1 to L16-2-3 segments. **a,c,e** PCR products were detected only in the targeted deletion mutants, yielding fragments of 5,753 bp (L16-2-1; a), 4,563 bp (L16-2-2; c), and 5,164 bp (L16-2-3; e). **b,d,f** Verification deletion of the internal region using three primer sets located within each targeted segment in the transformants shown in (a, c and e). PCR products of 1,259, 1,037, and 3,531 bp (L16-2-1; b); 3,531, 434, and 2,766 bp (L16-2-2; d); and 2,766, 1,567, and 2,464 bp (L16-2-3; f) were expected from primer sets 1, 2, and 3, respectively, in the presence of the intact wild-type locus, whereas no amplification was observed in the deletion mutants. M, molecular weight marker; W, wild-type.

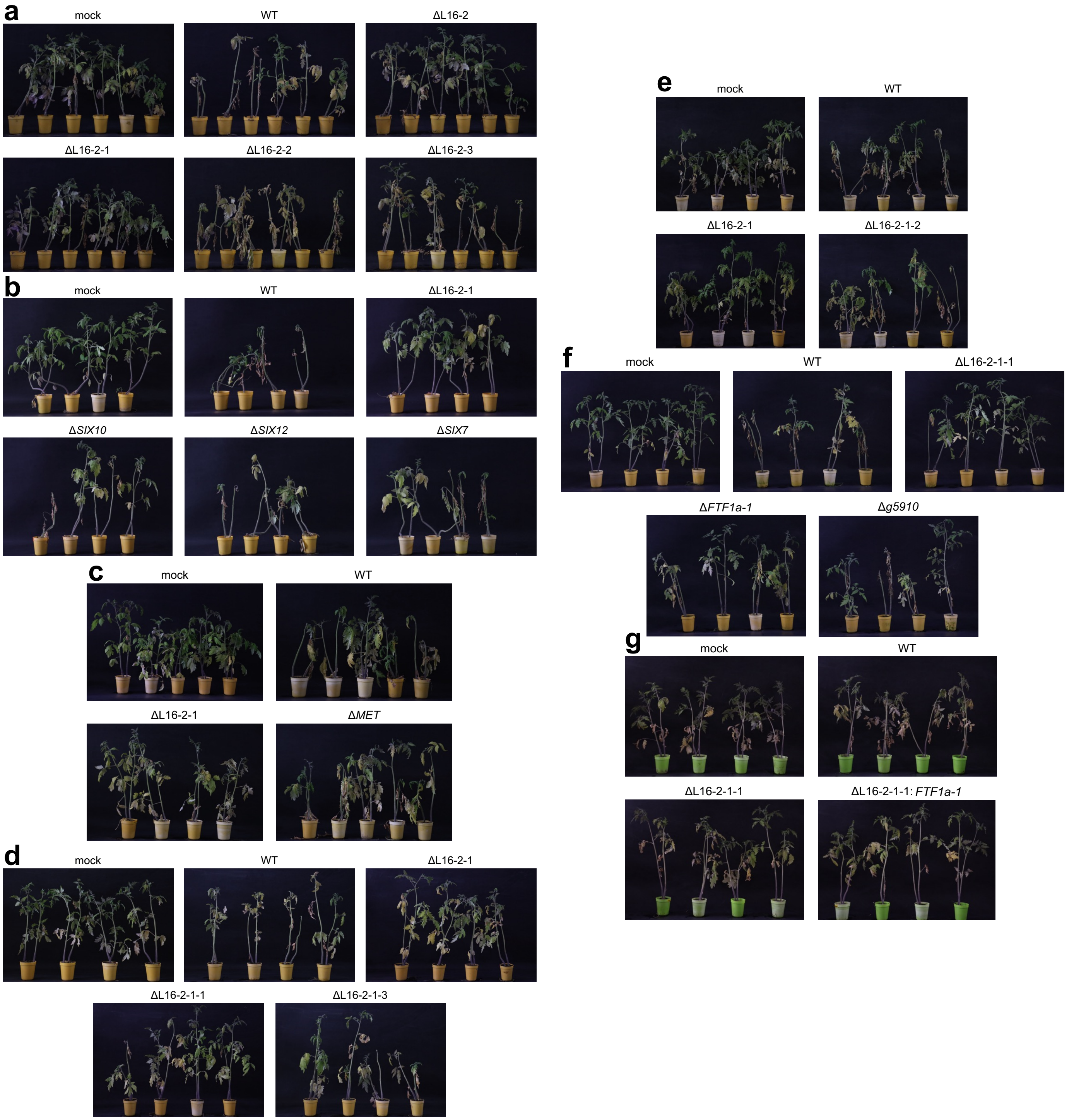


**Supplementary Figure 14: Representative external disease symptoms of tomato plants inoculated with mutants derived from the L16-2 segment. a–g** Representative images of tomato plants inoculated with deletion mutants derived from L16-2 (a), deletion mutants and gene disruption mutants derived from L16-2-1 (b–e), and gene disruption mutants and complemented transformants derived from L16-2-1-1 (f and g). For each mutant, disease symptoms were evaluated by comparison with mock- and wild-type-inoculated plants.

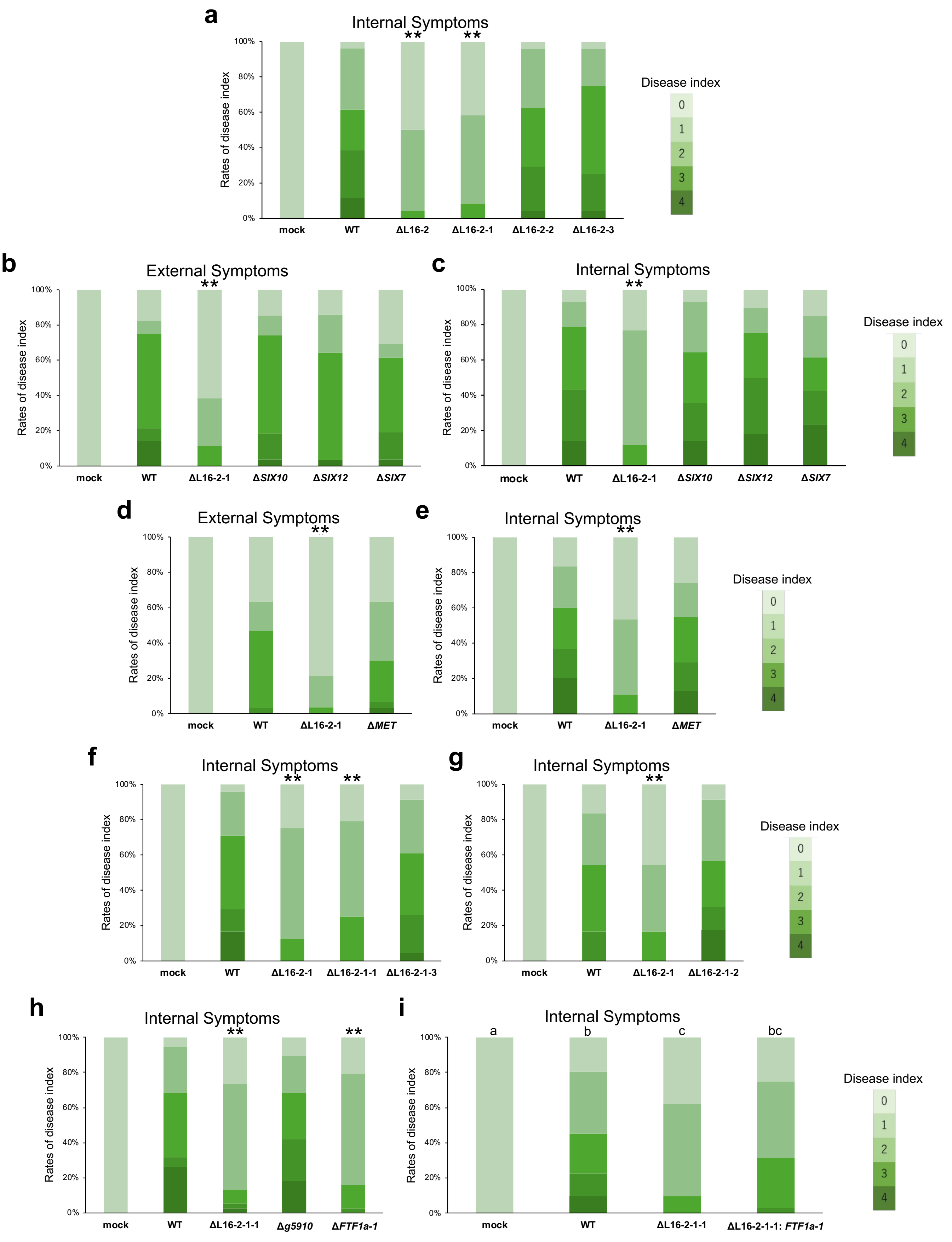


**Supplementary Figure 15: Virulence analysis of mutants derived from the L16-2 segment based on**

**internal and external disease symptoms. a–i** External disease symptom-based evaluation is shown in (b and d) and internal disease symptom-based evaluation is shown in (a, c and e–i) for the indicated mutants. Disease index was scored as described in the Methods. Asterisks in (a–h) indicate significant differences compared with the wild-type (n = 23–38 biological replicates; *adjusted $P < 0.05$, **adjusted $P < 0.01$; Mann–Whitney *U*-test). Different letters in (i) indicate significant differences among the wild-type and mutant strains (n = 29–32 biological replicates; adjusted $P < 0.05$; Kruskal–Wallis test followed by Dunn's multiple-comparison test).

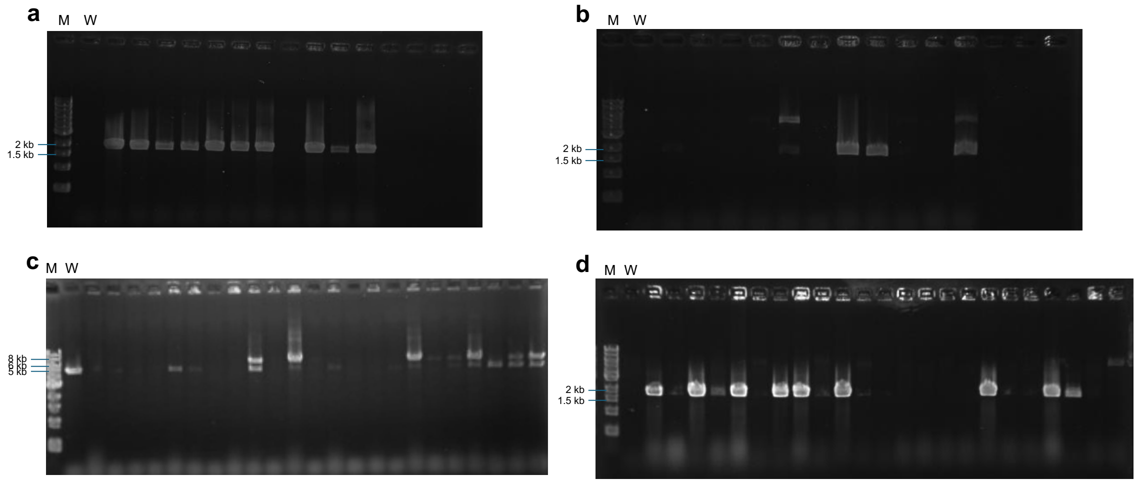


**Supplementary Figure 16: PCR-based validation of the single-gene disruptions within the L16-2-1 segment.** Representative agarose gel images confirming the targeted disruption of *SIX10*, *SIX12*, *SIX7*, and *MET*. **a,b,d** PCR products amplified using a primer located outside the FL region and a primer located within the *Hph* cassette were detected only in the corresponding disruption mutants, yielding fragments of 1,982 bp (*SIX10*; a), 1,977 bp (*SIX12*; b), and 1,972 bp (*MET*; d). **c** PCR products amplified using primers flanking the FL and FR regions yielded a 5,509-bp fragment in the wild-type and a 7,258-bp fragment in the *SIX7* disruption mutant. M, molecular weight marker; W, wild-type.

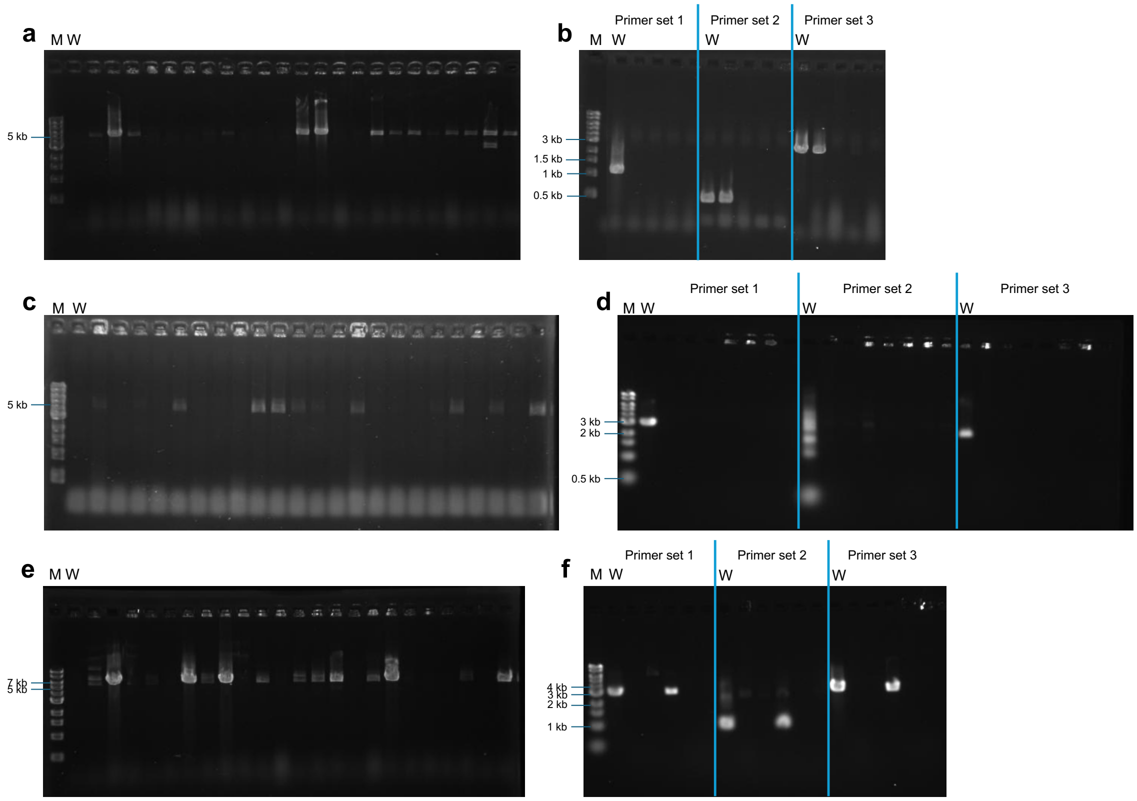


**Supplementary Figure 17: PCR-based validation of the large-scale deletion of the L16-2-1-1 to L16-2-1-3 segments.** Representative agarose gel images confirming the targeted deletion of the L16-2-1-1 to L16-2-1-3 segments. **a,c,e** PCR products were detected only in the targeted deletion mutants, yielding fragments of 5,269 bp (L16-2-1-1; a), 4,999 bp (L16-2-1-2; c), and 5,739 bp (L16-2-1-3; e). **b,d,f** Verification deletion of the internal region using three primer sets located within each targeted segment in the transformants evaluated in (a, c and e). PCR products of 1,259, 523, and 3,065 bp (L16-2-1-1; b); 3,065, 210, and 2,226 bp (L16-2-1-2; d); and 3,239, 1,037, and 3,531 bp (L16-2-1-3; f) were expected from primer sets 1, 2, and 3, respectively, in the presence of the intact wild-type locus, whereas no amplification was observed in the deletion mutants. M, molecular weight marker; W, wild-type.

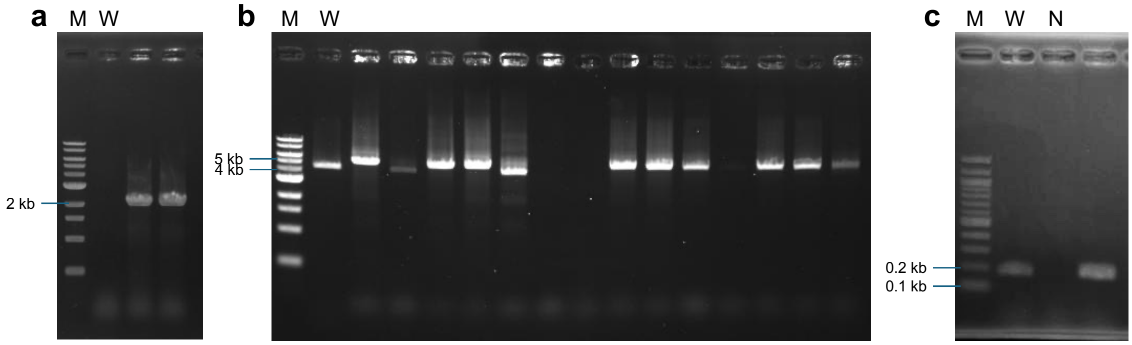


**Supplementary Figure 18: PCR-based validation of gene disruption and the *FTF1a-1* complemented strain.** Representative agarose gel images confirming targeted disruption of *FTF1a-1* and *Fol_MAFF103036_v2_g5910*, and complementation of *FTF1a-1* in the Δ L16-2-1-1 background. **a** PCR products amplified using a primer located outside the FL region and a primer located within the *Hph* cassette were detected only in the *FTF1a-1* disruption mutants, yielding a fragment of 2,067-bp. **b** PCR products amplified using primers flanking the FL and FR regions were detected in both the wild-type and the *Fol_MAFF103036_v2_g5910* disruption mutants, yielding 4,721-bp and 5,505-bp fragments, respectively. **c** PCR products amplified using *FTF1a-1*-specific primers were detected in the wild-type and the complemented transformant, yielding a 185-bp fragment. M, molecular weight marker; W, wild-type; N, negative control.

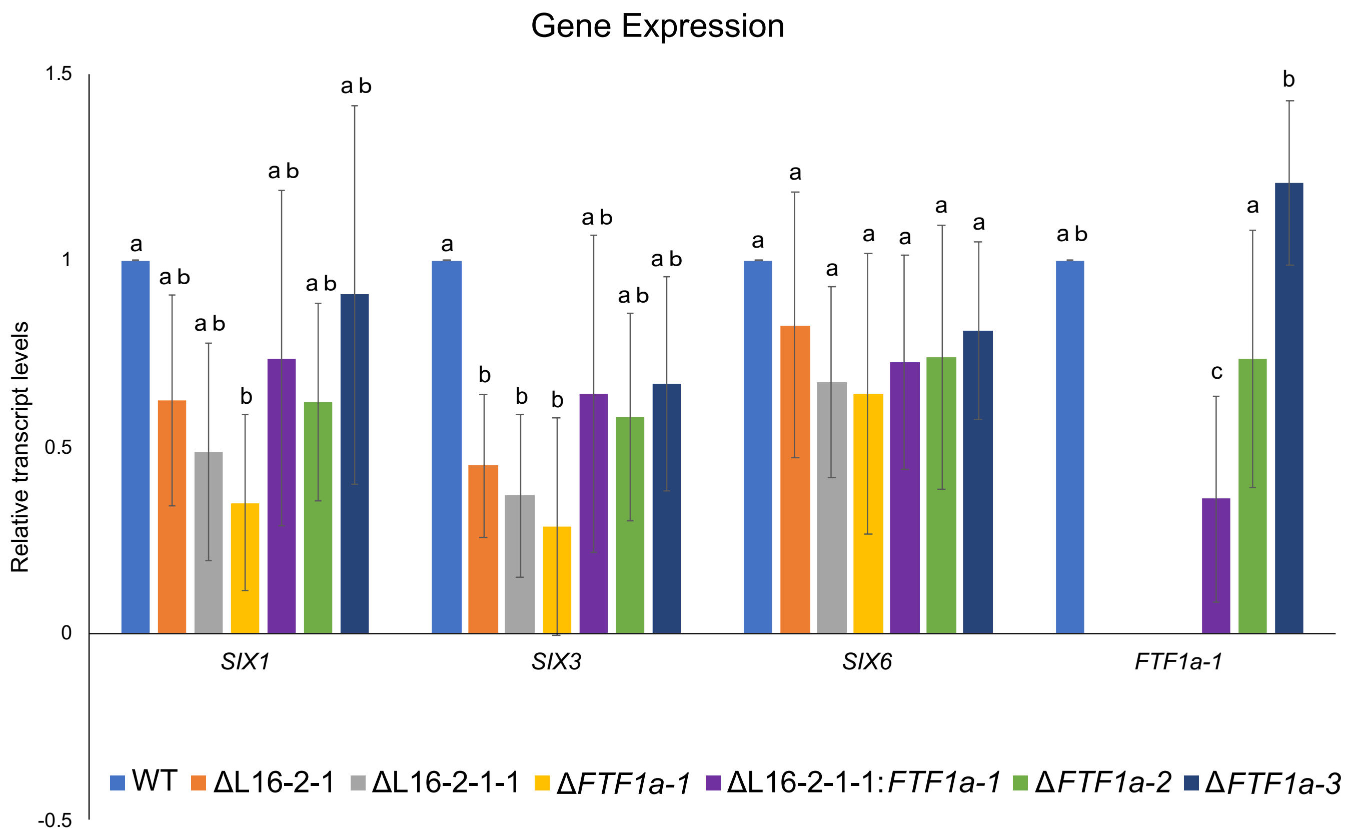


**Supplementary Figure 19: RT–qPCR analysis of gene expression in AC deletion mutants, gene disruption mutants, and complemented strains.** Relative transcript levels of the indicated genes were quantified by RT–qPCR using RNA extracted from tomato roots at 7 dpi. Expression levels were normalized to the reference gene *EF1α* and are shown relative to the wild-type strain. Error bars indicate standard deviations from three biological independent experiments. Statistical significance was assessed by one-way analysis of variance (ANOVA) followed by Tukey's multiple-comparison test. Different letters indicate significant differences among strains within each gene (adjusted $P < 0.05$).

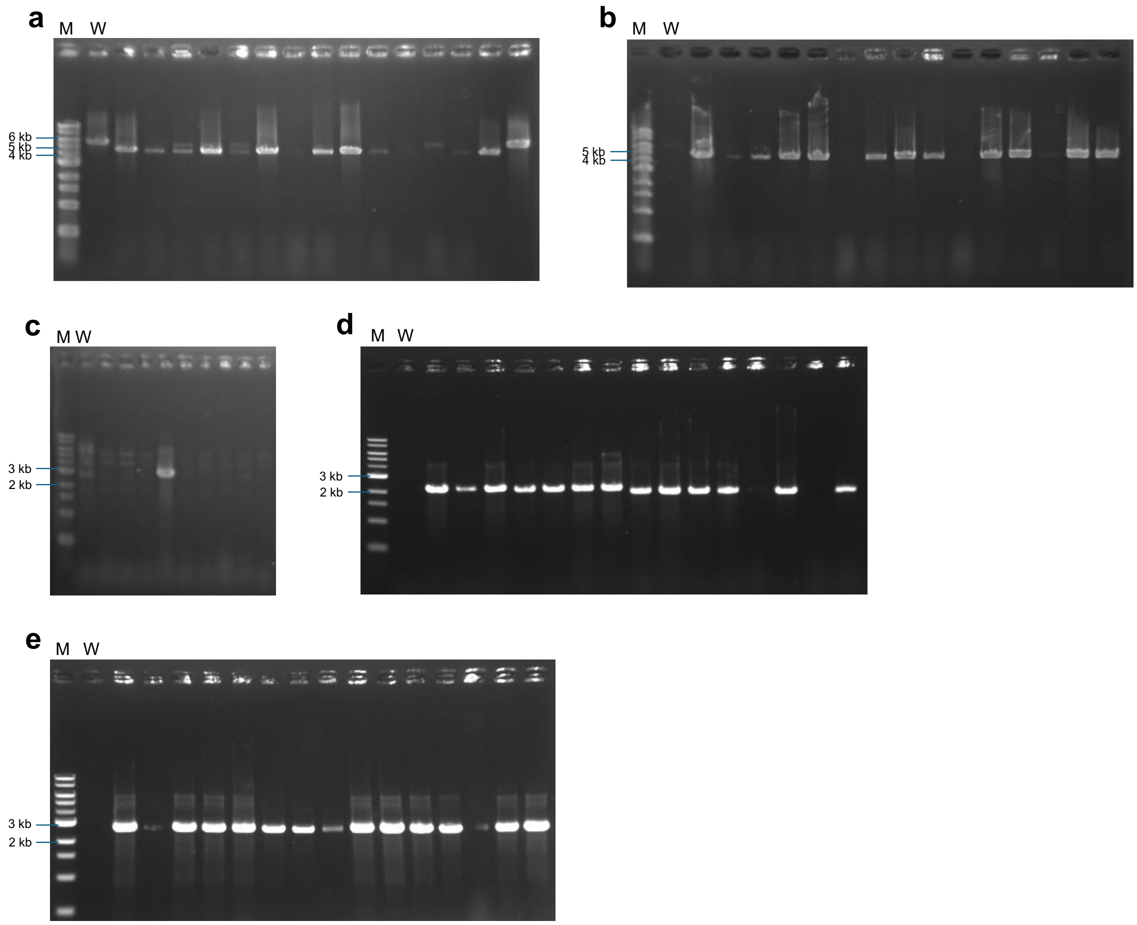


**Supplementary Figure 20: PCR-based validation of *FTF1* paralog disruption.** Representative agarose gel images confirming the targeted disruption of *FTF1a-2*, *FTF1a-3*, *FTF1c-1*, *FTF1c-2*, and *FTF1c-3*. **a,b** PCR products amplified using primers flanking the FL and FR regions were detected in both the wild-type and the *FTF1a-2* disruption mutants, yielding fragments of 5,732 bp and 4,695 bp for *FTF1a-2* (a), and 4,554-bp in the *FTF1a-3* disruption mutants (b). A 9,278-bp fragment was expected in the wild-type *FTF1a-3* locus but was not detected, likely because of its large size. **c–e** PCR products amplified using a primer located outside the FL region and a primer located within the *Hph* cassette were detected only in the corresponding disruption mutants, yielding fragments of 2,655 bp (*FTF1c-1*; c), 2,483 bp (*FTF1c-2*; d), and 2,402 bp (*FTF1c-3*; e). M, molecular weight marker; W, wild-type.

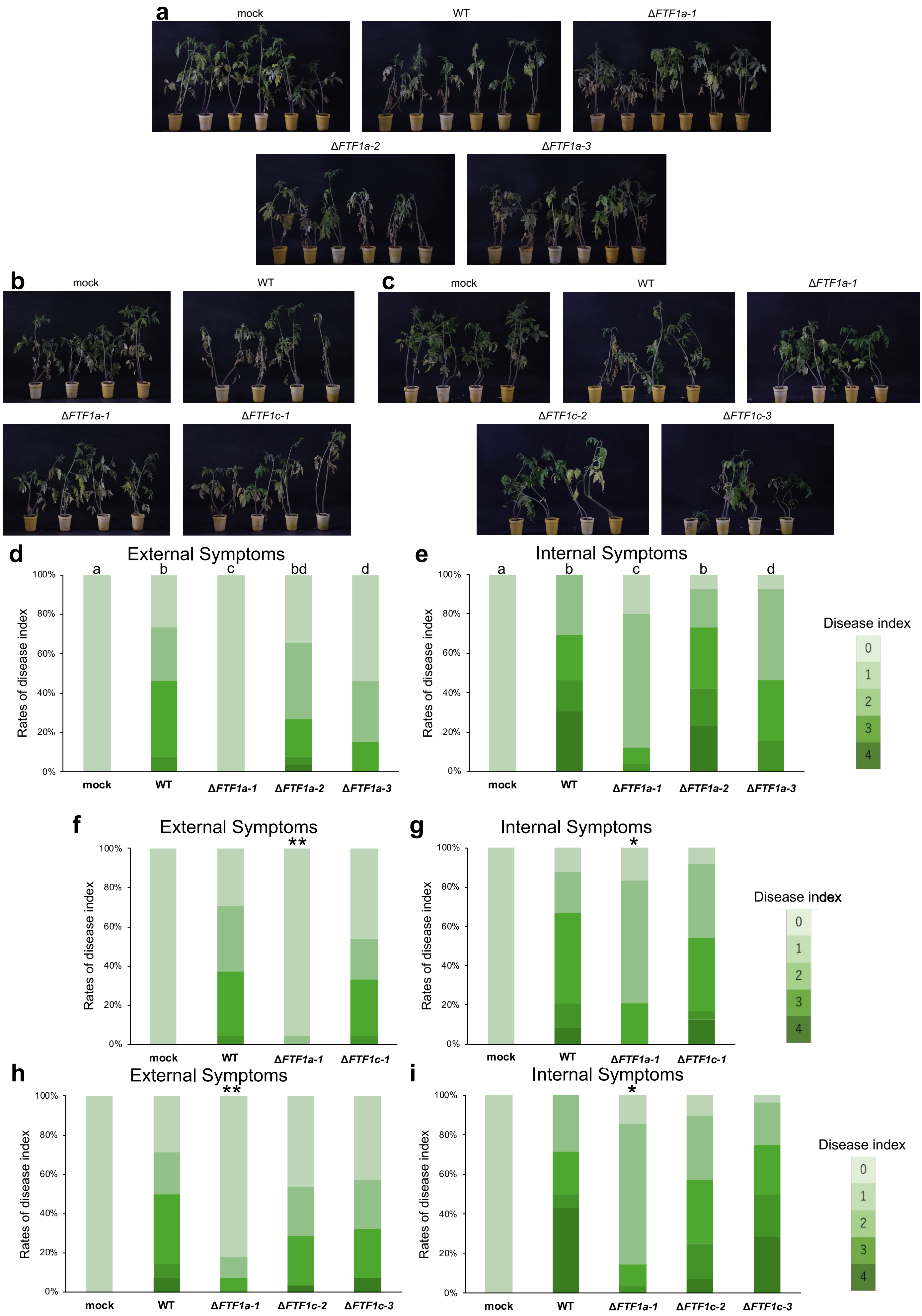

a
mock
WT
ΔFTF1a-1
ΔFTF1a-2
ΔFTF1a-3
b
mock
WT
ΔFTF1a-1
ΔFTF1c-1
c
mock
WT
ΔFTF1a-1
ΔFTF1c-2
ΔFTF1c-3
d
External Symptoms
a b c bd d
Rates of disease index
100% 80% 60% 40% 20% 0%
mock WT ΔFTF1a-1 ΔFTF1a-2 ΔFTF1a-3
e
Internal Symptoms
a b c b d
Rates of disease index
100% 80% 60% 40% 20% 0%
mock WT ΔFTF1a-1 ΔFTF1a-2 ΔFTF1a-3
Disease index
0 1 2 3 4
f
External Symptoms
**
Rates of disease index
100% 80% 60% 40% 20% 0%
mock WT ΔFTF1a-1 ΔFTF1c-1
g
Internal Symptoms
*
Rates of disease index
100% 80% 60% 40% 20% 0%
mock WT ΔFTF1a-1 ΔFTF1c-1
Disease index
0 1 2 3 4
h
External Symptoms
**
Rates of disease index
100% 80% 60% 40% 20% 0%
mock WT ΔFTF1a-1 ΔFTF1c-2 ΔFTF1c-3
i
Internal Symptoms
*
Rates of disease index
100% 80% 60% 40% 20% 0%
mock WT ΔFTF1a-1 ΔFTF1c-2 ΔFTF1c-3
Disease index
0 1 2 3 4

**Supplementary Figure 21: Representative external disease symptoms of tomato plants inoculated with *FTF1* disruption mutants and virulence analysis in MAFF 103036 based on internal and external disease symptoms. a–c** Representative images of tomato plants inoculated with gene disruption mutants of *FTF1* paralogs. For each mutant, disease symptoms were evaluated by comparison with mock- and wild-type-inoculated plants. **d–i** External symptom-based evaluation for (d, f and h) and internal symptom-based evaluation for (e, g and i) of the indicated *FTF1* paralog disruption mutants. Disease index was scored as described in the Methods. Different letters in (d and e) indicate significant differences among the wild-type and mutant strains (n = 25–26 biological replicates; adjusted $P < 0.05$; Kruskal–Wallis test followed by Dunn's multiple-comparison test). Asterisks in (f–i) indicate significant differences compared with the wild-type (n = 23–28 biological replicates; *adjusted $P < 0.05$, **adjusted $P < 0.01$; Mann–Whitney *U*-test).

```
Position 1-120
Ftf1a-1     MSGRAVLSPQHAQASFDSGLQLYGDVPELNAVPPPSHSALMDFTRFDDFAFAYYGLPDQSSLVSLVDHTHTFQSLTTFPQHQAISGLAHSGLPFGTLPTDNYNQSMEDSKAPPDRTSPAS
Ftf1a-2     ******SN*****Q*****P***R*****H***S********S*H********H***********A*****S**P*A******M***************G*RS****G*EP*********
Ftf1a-3     ***H**FE*STCPIVLRFRS***R*****H***S**********H***V****************L********P*A******M***************G*RS****G************
Ftf1c-1     ----------------------------------------****N************VRA**************P*AL*********************G*R*****G***H********
Ftf1c-2     ----------------------------------------****H**************A**A*******M***P*A******M*********L*****G*RS****G************
Ftf1c-3     ----------------------------------------****QL*********************Q******P*A******M***************G*RS****G************

Position 121-240
Ftf1a-1     NALEDPTTDEFGLASPNRAGGTDLGGKPKEDKADATPAWSELKTKAGKERKRLPLACIACRRKKIRCSGEKPACKQCLHSCIPCVYKVATRKAAPRTNCMAMLDKRPKRMEERAIKAISK
Ftf1a-2     ***************R***D*******H****V************************M*****************H**R*R*L*****T********GYI*****PL******I**V*P*
Ftf1a-3     **F*N**********C***D********R****G******G***********************V**********H****H*******T*********Y*V*******C****I***TP*
Ftf1c-1     E*P************RS****I*P*********************************V******S*********EH**R*Y*******T************************V*E****
Ftf1c-2     ***************R**V**I******************G*****************P***************EH**R*Y******IT********DY*********C****V****P*
Ftf1c-3     *****S*********R***D*****V**********A***G*R*******R***********************EH**C*Y*******T****G***DY**************V******

Position 241-360
Ftf1a-1     SDQEVASSVTHPVVKQAIPGTVTSSRPTKKRGAEEAFEPDLEAWAKASSEPKIEGDDGSSSLQVQEGEENKLQHEGTEALPSREIQEHLAEVFFENIYGQSYHLLHKPSYMRKLKNGTLP
Ftf1a-2     **********RA***PV*****P**K*********V*G***D*****P*K*******RP*C*****A***M*****A*****K***********D*************************
Ftf1a-3     *******F**R****P****A*P**K***********GL********P*K*******RP****A******************E***********D***************L*********
Ftf1c-1     **********R****P******P**Q***********G*********P****T************************D****K***********D*************************
Ftf1c-2     **********R****P******P**K*****S*****G*****C***P********N*P*N*********************KD******D***D*************************
Ftf1c-3     L*E*******C****S******P**K*****S*****G****T***VP*****************GK*G****Q********Q*******L**D**C***********************

Position 361-480
Ftf1a-1     PVLVLTVCAVAARFTSNPLVSSSGPEFLRGEEWASHARDICTRRYEWPNLTILTCLLILGLHEFGTCQGGRSWALGGQAIRMAFALQLHKDLEYDPSGRNGTTTQLSFIDREIRRRIMWA
Ftf1a-2     **************************L*********Q*******************************S********H*********************HS*K*****************
Ftf1a-3     **************************************E****Q****S***************************************************AK******************
Ftf1c-1     ****************S*******************************************************************************LD****K*****************
Ftf1c-2     ******************************************K***********************************************************K*****************
Ftf1c-3     ****************S***N**R**********************C****************************S**************************K*****************

Position 481-600
Ftf1a-1     CFLMDRFNSSGTDRPMFIREDTIQIPLPVKEKYFQFDMPAPTEMLDGRVPHPPSPNDGQIANSRENMGVAAFLIRAIALWGRIITYLSQGCKDLDPNPLWEDESHYMKHLDDVVNLEASL
Ftf1a-2     *************************************L********Q****V********DA****************************G*******M*******V***N*********
Ftf1a-3     ***L***************************************L***Q************DA***********************************L***E*P**I***N*A*******
Ftf1c-1     ****************V*******************GL**********************DV***************T*****T*********************H****N*A*******
Ftf1c-2     ************************************G***********************DV*********************T*C*********S*****GK****N**N*A*******
Ftf1c-3     *********FE************EL*****************************K*****DT****************************************K*******N*I*******

Position 601-720
Ftf1a-1     PLSLKHSAENLEVHKTENTASQFLFMHICLQHNILFVNRAAMSARKQHGVHDDFVSEASKRAFNAANRISELLREAEQSGCFVSAPFAGYCAFSSTTVHILGIISRNPFTKLAAQANLTT
Ftf1a-2     *****Y*******************************S*********R******F******T*D**************LR****************************SM**********
Ftf1a-3     ****NY************A****************L*S*********Q******F****Q*T*S*******F*******R***************A*********C**SM*PT*E*****
Ftf1c-1     *****Y*************P***************L*S****************FF*****T*********************************A***SV*******SM**********
Ftf1c-2     *****Y*************P***************L*S****************F******T***************************************A******SM**T*E*****
Ftf1c-3     *****Y***********K*P****L**L*********S*************V**F******T*****************************************M****SM**T*******

Position 721-840
Ftf1a-1     NIKYLHRMKKYWGMFHWMVENVRTQYRNVLDAMRAGANVEERATQPSFLQYGDWFNRYPRGLSDAEFMDPATHKRKDSGADGVLEAKRELRSVEEYFTLPTPRRVENKDTIRATAPKRKQ
Ftf1a-2     *V****K***************H*****A**********Q*****S**********C**H************L**************S**Q************SA********A******
Ftf1a-3     *V****K************************H*****V********L*******Y********************************P********S****************A******
Ftf1c-1     *V****K********************Q*****************L*****************H**********************NP**************Q********S*A******
Ftf1c-2     *V****K**************************************L**************************************D**P******K******************A******
Ftf1c-3     *V****K*******************************************************F************************P*************************A******

Position 841-960
Ftf1a-1     SAKKQAGMPAQPGQHLDSLQSIDADAVSQERKFSGGLGLQITGAAGFNPLAASNQQSPDFSTTISPTRPANMTPFAHHAHTPTFFPPELLAMNFGQGSNGNIDPLDRQLIYGGYSMDAST
Ftf1a-2     *****T***************T******************T****D********P*N*A****MP**S*****A*****Y*****************************V**********
Ftf1a-3     *****T***************T****************************S*****N******M***S*******S*Y************SID***************************
Ftf1c-1     N****TD****SDRN**W***T**********L**********S******T************ML*MS*V***S*************QPF******************F***********
Ftf1c-2     *****T************************H*************************N******M***S*****A***Y******************D***********************
Ftf1c-3     *****T**************I*EP******H************V*********KT*I******M**MS********Q****H*******************T************L*****

Position 961-1079
Ftf1a-1     GLGGGQDMMSGLGWDTVALGAQPDGRLQSRPSNAKAGMHGQSAGMADGAGLSRPEASSAWFMPFNMEPPDMDQDAGFNMGGIDPFTGVFGGGGSGLATPNALGGLILGHCRGSSTHSTR
Ftf1a-2     ***D****T***D*****S******G**G*R**V*****************GL****************IG**P**********A********************QQ*P----------
Ftf1a-3     **C***************S******C**G**P*V*****************G**************A*****NP******F******L*****************QQ**----------
Ftf1c-1     S**D*--------------------------------------------------------------------------------------------------------------*TWA
Ftf1c-2     ***H*--------------------------------------------------------------------------------------------------------------*TWS
Ftf1c-3     ***D*--------------------------------------------------------------------------------------------------------------YTWA
```

**Supplementary Figure 22: Multiple sequence alignment of amino acid sequences of Ftf1 in MAFF 103036.** Alignment of the amino acid sequences of Ftf1a-1, Ftf1a-2, Ftf1a-3, Ftf1c-1, Ftf1c-2, and Ftf1c-3. Asterisks (*) indicate amino acid residues identical to those of Ftf1a-1, whereas amino acid substitutions are explicitly shown. Gaps introduced to optimize the alignment are indicated by dashes (–). The predicted zinc-finger-like DNA-binding region (amino acid residues 175–235 of Ftf1a-1) is highlighted in blue.

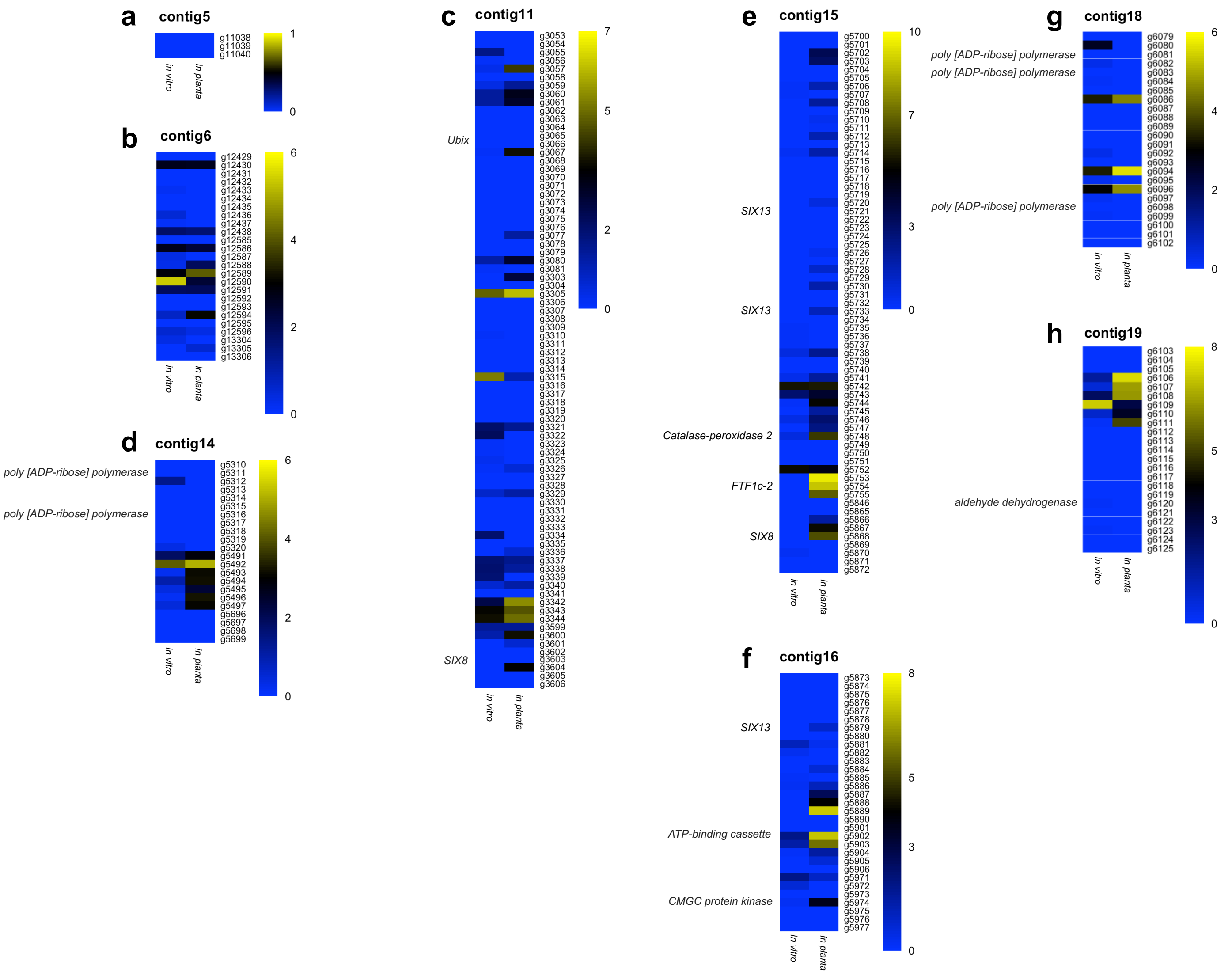


**Supplementary Figure 23: Transcriptome analysis of genes within accessory chromosome regions not covered by the deletion library in MAFF 103036. a–h** Heat maps show the expression profiles of genes within contig 5 (a), contig 6 (b), contig 11 (c), contig 14 (d), contig 15 (e), contig 16 (f), contig18 (g), and contig19 (h) under *in vitro* and *in planta* conditions. Each row represents a gene, and each column represents a condition. Displayed values represent $log_2$-transformed TPM. The color scale indicates relative expression levels, with blue representing low expression and yellow representing high expression. Gene names are indicated on the right side of each heatmap. Only genes showing high sequence similarity to those of 4287 are included. Genes upregulated under *in planta* conditions compared with *in vitro* conditions were defined as candidate virulence-related genes.

Position 1-180
*FTF1a-1* --------------------------------------------------------------------------------------------------------------------------------------------------------------CAAACTTATTGCATCCCACTGCAGG----------GAAGTGTTAAAGGCAGCAGGATAT
*FTF1a-2* TACAGTATGAGGAAGATCACGATGCTTGGCAGAGCCCATACAAGAGACCAGACCACGAGCTAAGCCGAACGAGCCGCAACAGCTACGCGTTCTTGCATCATTTCAGCCTCCGCAACGGTTC**C***CT**T*CG**A***C*TTC---------------------------------
*FTF1a-3* --------------------------------------------------------------------------------------------------------------------------------------------------------------ATGC***T****CC*ATG**CATCCTACATGTAGG******G*G************G*
*FTF1c-1* -----------------------------------------------------------------------------------------------CTAGCGCAGCT----------AACACAACTTACACACCTG-CGA*T****T****CC***C*CC*CCC-----------------------------------------
*FTF1c-2* ------------------------------------------------------------------------------------------------------------------------------------------------------------------------------------------------------
*FTF1c-3* -----------------------------------------------------------------------------------------------TAGAGAAGCTAGGGCCTATCAACGCGACTTACACACCTG-CGA*T****T***-----------TCC-----------------------------------

Position 181-360
*FTF1a-1* TCATACATCAATCATAATATTGCAACCAGACTTGTAAATCGTATGATAGAGTCTCTAGCCTTG----------------TAGGCTTGGTACGTACAGTACAGTCCCGTTTTGGGTGATGTGTGATGTCAGGCGCAGACTGTTCCCAGCTTCAGCGCGGGGAGCCACCGACTACATACGTA
*FTF1a-2* ----------------------------------------------------------------------------------------------------------------------------*CCCC*GCCGG************C*******C***T************T*************
*FTF1a-3* ******GC*C***************T******C**********C*G************T*C----------------C***G*********T**********T*****C********C*******G*******C******CC***T************T******-------
*FTF1c-1* -------C***CTG**CA*CA*****ATAGA*CTA-**CA*A*AA*A**C**T****TGT*ATTAATGCATGTAAACAAGCA*T*A*AATATACA**ACG**GAATTC*GATT***G*C*TAA*AAC*TA*T**T**T*-------------------------------------------
*FTF1c-2* ------------GG****GC*TATGAATTTGACTAT**GAA**ACC*T*CC***T**ATTCACATATTTTTGACCGTAA*GA*GGC*TA*GT*C*TTGC*T*GAGGCA*GGT*AG*CGT*A*C*G*CGTT*T*AT*TG***----------**A*GA**A**AAGAA*CGGCATC*AAA*
*FTF1c-3* -------C*C*CGG**CA*CA*A***ATAGA*CTAG**CA*A*AA*A**CA*T****TGT*G---ATGTGTATAAATAACTA*T*AAAATATACA**ACG***AATT**GGTT***G*C*TAA*AAC*T**TCTT**T*C-------------------------------------------

Position 361-540
*FTF1a-1* CCTGTTGCTTGCTTCTCCATAAACTTCCCGGACGGCTTCAGGTCACCTCAACCTCAACCTCAACGCTCTTGCGCTGCAACTTTTTAAACTGCACAACAGCTGTACCAGCCAACGTC--------------------------TTATGAAGGCTTTGTTTTGCCCTGTCGGACTGCATTCG
*FTF1a-2* **C**********************CA*******A*************************************A*A**********************************G***C**-----------------------TTAT****A******************A******TG*A*****
*FTF1a-3* -*******C********T*******C***A*********G****G******************************************************T*********CC*-----------------------******A***A*****-**C***TA*********G****
*FTF1c-1* --------------------------*G*CCTG*AG*G****C**GTGTG*GGC**C*ACA*CGC***G*ATT*AAT*T*A**---G*ACA*TTGC*GC*CAA*TT*C*G***CGTGCCTCGGCAGCAAAGGTCAAAGTTTTGCTAA***AAAC*A*G*A*GG*A*C***G*C-------
*FTF1c-2* *GC*G*C**GATCGGGTT*GCCCGGG**AA**TAC**GG**A*GTGT****A**T*TTGCACC*CACTG*ATC*AAT*T*A**GA*G*A*A*TTGC*GC*C*A*TT*C*GT*ACGTGCCTCGGCAGCAAAGGTCAAAGTTTTGCTAA***AAAC*A*G*CAGG*A*C***G*CC------
*FTF1c-3* --------------------------***CCTG*AG*G**A*C**GTGTG*GGC**C*A*A*CGC***G*ATT*AAT*T*A**GA*G*A*A*TTGC*GC*C*A*TT*C*G***CGTGCCTCGGCAGCAAAGGTCAAAGTTTTGCTAA***AAAC*A*G*A*GG*A*C***G*CC------

Position 541-720
*FTF1a-1* CAGCCACCAAACCCCTGTCTAACTCAGTCGCTTCGCTCCACCATCATCACCACAAGGGTCCGGTATATTTGTTTTCCATCGCCAGATTCTCGGCCTCGGGTGGCC-ACGTCGTAACTCTGTCTCTGGCAGAGCCATGTCCGGCCGTGCCG-TTTTGAGCCCTCAACATGCCCAAGCGTCC
*FTF1a-2* ****************C*******C*******************G**********GCCC*******************C**********************C*************-***********************************************.***C**A****************A*****
*FTF1a-3* ****************CG*****G************G*****G*****T**GCCC**********************G*******************-G*********T*********************************A*****T******A****************TA****
*FTF1c-1* ---------------------------------------------------------------------G*CCG****G**TATCCG******************-G**************------------------***********-******AT******T********CA****
*FTF1c-2* ---------------------------------------------------------------------G*CCG****G**TATTCG******************-GG*************------------------***********-******A****************CA**T*
*FTF1c-3* ---------------------------------------------------------------------G*CCG****G**TATCCG***C*********A*****-G**********C***-----------------**A*A***T**-******A*****G***********A****

Position 721-900
*FTF1a-1* TTTGATTCTGGTCTTCAACTCTATGGCGACGTGCCTGAATTGAACGCTGTGCCTCCTCCCTCTCACAGCGCCTTAATGGATTTCACACGCTTCGACGATTTTGCGTTTGCTTACTATGGTCTTCCTGACCAGTCTTCTCTCGTTTCCCTAGTAGATCACACCCACACGTTCCAATCTCTC
*FTF1a-2* **C*****C****CC********C********T**********C*************T****************************C*C*****A***************************C***********************A*********C*****************CA*******C*
*FTF1a-3* **C*****C****C**********G**********************C*******************T*****A***************************************G************C****************************************************C*
*FTF1c-1* **C*****C****C*********C*******T********C*******************A***************T*C*********AA***************************************T**G*G*********************************************C*
*FTF1c-2* **C*****C****TCC*******C*******T********C*****************************************A*************************************************G*********C*******************************T****C*C*
*FTF1c-3* **C*****C***AC-*********C*******A*******CTC*G*********TT**A****************************AG**G**********************************************************G*****A*******************C*

Position 901-1080
*FTF1a-1* ACTACGTTTCCCCAGCACCAGGCCATATCTGGCCTTGCACATAGCGGTCTGCCGTTCGGCACCTTGCCTACGGACAACTACAACCAGAGCATGGAAGACTCCAAA---GCTCCCCCAGATCGGACATCTCCCGCATCCAACGCCCTCGAAGACCCGACTACCGATGAGTTTGGTTTGGCT
*FTF1a-2* ***G*********************G*****************************************************G****CG**G**************G****G**CTCC*C*****C*****************************************A***************************
*FTF1a-3* ***G*********************G**********************************T**********G****CG**G*******T*******G*******---**C**G*T******************************T*****A***************************
*FTF1c-1* ***G**C******************T*****************************************************G****CG***************G*****G---**C*A**************************G*G****C************************************
*FTF1c-2* ***G*********************G******************************C***********************G****CGA*G***************G*******---**C*****************************************************************
*FTF1c-3* ***G*******T*************G*****************************************************G****CG**G******T*******G*******---**C**A**************C*****************************T****************************

Position 1081-1260
*FTF1a-1* TCCCCTAACCGTGCGGGTGGCACTGATCTAGGCGGTAAACCTAAGGAGGATAAAGCCGATGCCACACCTGCGTGGAGTGAACTTAAAACAAAGGCTGGCAAGGAAAGAAAACGCCTCCCGCTTGCTTGCATTGCTTGTCGCCGAAAGAAGATCCGTTGTTCTGGCGAGAAACCCGCCTGC
*FTF1a-2* ****G**********A******A***********C****A*************************T*****************************************************T********C*******G**A**C*************************A********************
*FTF1a-3* ***TG*********A*A******A**********G*****G**************G************************G*******G*****************************************A**C***************G**********A**************************
*FTF1c-1* ****G**G*************TA****C**T*****************************************************G**********T***********************************G*****A*****************G******C**A**************************
*FTF1c-2* ****G*********T******T*TA*************************************************************G*******G****************************************C*A****************************A***************************
*FTF1c-3* ****G**********A******A*****************************************************************G*******GG*****************************G***************************************A**************************

Position 1261-1440
*FTF1a-1* AAGCAGTGCCTACACTCATGTATCCCATGTGTCTACAAGGTTGCGACTCGGAAGGCTGCGCCTCGGACAAATTGCATGGCTATGCTCGATAAGCGACCGAAGCGCATGGAAGAACGCGCCATCAAGGCCATATCCAAGTCGGATCAGGAAGTCGCCTCATCTGTGACTCACCCCGTGGTC
*FTF1a-2* *****C**T****G****C******T************A*A**********************GG**A***A***************C**T*****************AT********T***C***********************A*******T****A****G*G*********
*FTF1a-3* *****C*********CA***********************A*****A**********************A*****G**************************T************************T**************************A*******T*A*****G********
*FTF1c-1* G****C**T****G****A**************T*******A***************************************************T***************T****G*********A*****************G*****C**A*****G**********
*FTF1c-2* G****C**T****G*****AC**************T******A**A**********************G***A***************************************T**************G*T*****************C*******C****************A*********A****G**T******
*FTF1c-3* G****C**T****TG*****A**************T**A**A**************G**********G***A***************************A**T**************************T****G****************A*********A***TG*********

Position 1441-1620
*FTF1a-1* AAACAGGCGATACCAGGAACCGTAACTTCCAGTAGGCCAACCAAGAAGCGCGGCGCCGAAGAAGCATTTGAGCCTGATCTGGAAGCTTGGGCGAAGGCATCTTCGGAGCCAAAGATTGAGGGCGATGATGGATCCAGCAGCTTACAAGTCCAGGAAGGGGAGGAGAATAAGCTGCAACAT
*FTF1a-2* ****C**T***********T***C*********A************************G****T***C*G*************T*************GC*****A***********************A*GC*****T****G*************C***********T*T*******
*FTF1a-3* ****C**********G*T***C*********A****************************G*********C*G**T**********************GC*****A*************************A*GC*******************G***C******************************
*FTF1c-1* ****C*************T***C*********CA***************************G*********C*G*****************************GC************C***************A*G********G******************************C
*FTF1c-2* ****C*************T***C*********A*****************A********G*********C*G********************C*******GC*****************************A***GC******A*********T*****************************
*FTF1c-3* ***TC*************T***C*********A******************A*******G*********C*G**********GA**********TGC******************************G***********G*****T****G*AA*****GC**C**********A

Position 1621-1800
*FTF1a-1* GAAGGCACCGAAGCACTCCCCTCCAGGGAGATACAGGAGCATCTGGCAGAGGTGTTTTTCGAAAACATCTACGGTCAATCCTACCATCTTCTACACAAGCCAAGCTATATGCGAAAGCTAAAGTGAGTTCCTTTTGGTTGA-TCTCACGTATCCTGCTAACTGTCAACAGAAATGGCAC
*FTF1a-2* *****G**********A*****************************C*********T***********T***************************************************T****T*****A*****************************T**
*FTF1a-3* ***********************GA************************C****************C*********T*********T*********************C*******T*****A****GT***************A****T******T****G***************************
*FTF1c-1* **********C**********A*********************************C*********T**********************************************T*****************A*************G*********************
*FTF1c-2* **********************A***C*****A****************T*********C*****C*****T*****************************************T*********C*A*************************************
*FTF1c-3* ***********************CA*********A************C********C*******GT**C*****T*********T***************************T*****T*****A******C*T*****************************

Position 1801-1980
*FTF1a-1* TCTACCTCCGGTGCTTGTTCTCACAGTGTGCGCTGTAGCTGCTCGTTTTACCTCAAACCCTCTAGTGAGTTCTTCAGGGCCTGAATTCTTACGCGGTGAAGAATGGGCATCACACGCTCGAGATATTTGCACCAGACGATACGAATGGCCAAACCTCACCATCTTGACGTGTCTTCTCAT
*FTF1a-2* A****************************************************CT***********************C*********************************A************************************************************
*FTF1a-3* A**************************************************T*********C*****************************************G**************A*************G**************A*************************
*FTF1c-1* A********************************************************G*********************************************************************************************C****A************
*FTF1c-2* A*********************************************************************************************************************A*****************************A**********************
*FTF1c-3* A***************************************************T**G*G*********A*A*******A***********************************************************T****************A***********

Position 1981-2160
*FTF1a-1* TTTGGGCCTTCATGAATTTGGAACGTGCCAGGGCGGCCGTAGCTGGGCCCTGGGTGGACAAGCTATCCGAATGGCTTTCGCTCTCCAGTTACATAAAGACTTGGAATACGATCCTCGGGCCGTAATGGCACCACAACGCAGCTCAGCTTCATTGATCGAGAGATTCGGCGACGCATAAT
*FTF1a-2* ********************************A*****************************T*****T*************************************************C**A****A*************************************************
*FTF1a-3* *******************************************C**T*********A****G***************************G**************************************G***A**********************************************
*FTF1c-1* *******************************************************************************************************************T**A*****A**A***********************************************
*FTF1c-2* *******************************************************************************************T*******************************A*****************************************************
*FTF1c-3* ******************************************************A*C**G************************************************************A*****************************************************

Position 2161-2340
*FTF1a-1* GTGGGCCTGCTTTCTCATGGATCGCTTCAACTCTTCTGGGACAGACCGACCCATGTTCATCAGGGAGGATACAATTCAGATTCCTCTGCCGGTAAAGGAAAAGTACTTCCAGTTCGACATGCCTGCGCCCACCGAGATGTTGGACGGTCGAGTACCTCATCCGCCGTCGCCCAACGACGG
*FTF1a-2* *****************************************T*************************************************************A*******C*******************A*************GT*******************
*FTF1a-3* *************T***************************T***********************************************T**A*******************G******C**********A*********************************
*FTF1c-1* *******************************************T*************G******************************T*****A***G*C**********************T*************************************************
*FTF1c-2* *****************************************A*****T*************************************************A***G*********************************************************************
*FTF1c-3* ***********************************T**A*****T************************G**C**********************A************************************G**********************A*****

Position 2341-2520
*FTF1a-1* ACAAATCGCGAATTCACGAGAGAACATGGGAGTTGCGGCCTTCCTGATTCGAGCCATTGCCTTATGGGGACGAATCATCACCTACCTGAGCCAAGGGTGTAAGGATCTAGACCCCAATCCATTGTGGGAAGACGAGTCTCACTACATGAAGCACCTCGATGATGTTGTAAACCTTGAAGC
*FTF1a-2* *********TG**G*********************************************************************************G*********************************************G*********T****A*****************
*FTF1a-3* *********TG**G************************************************G*************************************************T***********A**AC*********A*****T***A*******C*****************
*FTF1c-1* *********TG**GT*************************T******************G*******A*************G****C*****************************************************************C******T***A******C****************
*FTF1c-2* *********TG**GT*************************T**************************************G****C*****G*************************T*******C*********G*A****************C**T***A******C****************
*FTF1c-3* ***G*****TG**A*G********************************************************G*******T*****************************************A*******************T***A******A*************

Position 2521-2700
*FTF1a-1* TAGTCTGCCCTTGTCACTCAAGCACTCTGCAGAGAACCTGGAGGTCCACAAGACAGAGAACACGGCAAGCCAGTTTCTTTTCATGCATATCTGCCTGCAGCATAACATTCTCTTTGTCAACCGAGCTGCTATGTCAGCACGAAAGCAACATGGTGTACATGATGATTTCGTCTCTGAAGC
*FTF1a-2* *********************T****************C******************************T******************C*************************G*GT*************************G***********C*******T************
*FTF1a-3* *********************CT*************C*****************G**********A***************************************C*TC***G*T**C*C***************G**G*G*******T************
*FTF1c-1* *********************T*T*************C*********************C**************************************************A*G*GT*************************************T**************
*FTF1c-2* *********************T****************C********************C******************T**************************C**G*GT******************************************T***C*****
*FTF1c-3* *********************T****************C********************A***C*********C*********C*******A***********************G*GT*******************************T****************

Position 2701-2880
*FTF1a-1* AAGCAAGAGGGCCTTCAACGCTGCGAACCGAATATCCGAGCTTCTTCGTGAGGCTGAACAGTCGGGATGCTTTGTTTCGGCCCCGTTTGCTGGATACTGCGCCTTTTCCTCGACAACAGTTCATATCTTGGGTATTATCTCTCGCAATCCCTTCACGAAGCTAGCAGCCCAGGCCAATTT
*FTF1a-2* *********A*****G*****************************************T*C*****************************************************C*********************C**T*****************************
*FTF1a-3* G**TC*****A*A****G***********G*****T**AT*C*********C*********C*G**************A**C*C********************G********C*******************T*******A*C**T*****C*A*****G**********
*FTF1c-1* *********A***************************************************************************************************G**********C*G*G*************A*C*T*****************
*FTF1c-2* *********A*******A**********************************C*********************T******************************************C***********************A*C**T*******G***********
*FTF1c-3* *********A****************************C************************T*****************************************C********G*********C**T******GA*************

Position 2881-3060
*FTF1a-1* GACCACCAATATAAAGTATCTTCACAGAATGAAGAAGTATTGGGGCATGTTCCACTGGATGGTGGAGAACGTTCGCACTCAGTATCGAAATGTCTTGGACGCTATGAGAGCTGGTGCGAATGTCGAAGAACGAGCCACACAGCCGTCTTTCCTGCAATACGGAGACTGGTTAACCGTTA
*FTF1a-2* *********G*C*****C*****A*****************A**********************A******************C*************************T***********************T*A*****T*********************
*FTF1a-3* *********G*C*****C*****A****************T***************************************C*****************T***********************TA**C*****T*************A**************
*FTF1c-1* *********G*C*************A************************T**********************A************************************************************TT********T*******************
*FTF1c-2* *********G*G*************A************************T*********************************************************************************TT********T*******************
*FTF1c-3* *********G*C*************A************************T************************************C*****************************************T*********T*******************

Position 3061-3240
*FTF1a-1* CCCTCGCGGTCTTTCTGACGCTGAGTTCATGGATCCTGCCACTCACAAACGAAAGGATTCAGGAGCAGCGGCGTCCTCGAAGCGAAGCGCGAACTGCGATCAGTGGAGGAATA---CTTCACGCTTCCAACACCACGAAGGGTTGAGAATAAGGATACTATCCGCGCACAGCGCCGAA
*FTF1a-2* *****A*************************C*********T****************************************TC********A***************CTT**C***********************T*CC*****************************G**********
*FTF1a-3* ****************************************************************T*******************C*******************---**C************************************************G**********
*FTF1c-1* ***************C***********T*****C***************************************************T*C******************---*****************G*A****************************A****GG**********
*FTF1c-2* *************************************************************************C*******C**************A****---**********************************************************G**********
*FTF1c-3* **************T******************C**********************************************C*****************************---************************C**************G*************G**********

Position 3241-3420
*FTF1a-1* ACGAAAGCAAAGCGCCAAGAAACAGGCTGGCATGCCAGCACAACCTGGCCAGCATCTCGATTCGTTGCAGAGTATAGACGCAGACGCAGTCTCCCAGGAACGCAAGTTCTCGGGTGGTTTGGGATTACAAATCACAGGTGCAGCAGGCTTCAACCCTCTGGCAGCATCAAACCAGCAGAG
*FTF1a-2* *************************A*****************************************C*****************************************************G***CT**C*************A*************CT***********C*****A
*FTF1a-3* **********************G***A***A****************************C***T*************************************************G************T*******************************CT************A
*FTF1c-1* ************A*************A**A*********GT***A**G*A***************G***************C*****************C*A***************G************T*******************CA****************
*FTF1c-2* *************************A*************************************C***T**************************************************************************************************
*FTF1c-3* ******A******************A**************************T*******GC**********************A***********************************T*****************C***********AAC*****T

Position 3421-3600
*FTF1a-1* CCCGGATTTCAGCACCACCATATCACCTACGAGACCGGCCAACATGACTCCATTTGCTCACCACGCACACACGCCCACCTTTTTCCCACCCGAGCTGCTCGCAATGAACTTTGGGCAGGCTCGAATGGAAACATCGACCCGCTTGATCGTCAGCTTATCTATGGCGGATACTCAATGGA
*FTF1a-2* *****C************GC**********C*******************G************T****************T************************************************T***********G***********************
*FTF1a-3* ******************G*********C*********************T*****T*****************************AG*****************************T*****************************************
*FTF1c-1* *********T**********G*T*****T***C****TT**T*****T*****************T******GC***C*T**************************G************************AT********************
*FTF1c-2* ******************G*********C****************G*********T***************************************A*******************************A************************************
*FTF1c-3* ******************G*T*****T************************G****T***A***A******************************************C******************************************T

Position 3601-3780
*FTF1a-1* TGCTAGTACCGGCCTGGGTGGTGGCCAGGATATGATGAGCGGCCTTGGTTGGGATACCGTCGCTTTAGGCGCTCAGCCAGATGGACGCTTGCAAAGTCGGCCGTCAAATGCCAAGGCAGGCATGCATGGACAAAGTGCAGGCATGGCTGATGGGGCAGGATTGAGCAGACCGGAAGCATC
*FTF1a-2* ****************C*A*********C*********G*A******T*******C*************G*******G*******G******G***G****T*********************G*********T*****************C*A***G***T********
*FTF1a-3* ***********T***T*****************C***********T***********A**********C**T**************************************T********************************A***G**********
*FTF1c-1* *********A*T*******A*****TAC*TG*GCATTGTAG------------------------------------------------------------------------------------------------------------------------------------
*FTF1c-2* **************A***CAC*****TAC*TG*TCATTGTAG-----------------------------------------------------------------------------------------------------------------------------------
*FTF1c-3* ****************A***T*TAC*TG*GCATTGTAG------------------------------------------------------------------------------------------------------------------------------------

Position 3781-3960
*FTF1a-1* ATCCGCATGGTTCATGCCATTCAACATGGAGCTCCAGATATGGATCAAGATGCCGGCTTCAACATGGGCGGGATTGATCCGTTCACGGGAGTGTTTGGCGGAGGAGGCAGTGGTTTGGCAACGCCGAATGCGCTAGGTGGGCTCATACTTGGGCATTGTAGAGGGAGCTCTACCCACTC
*FTF1a-2* ***T****************************************A*G*******C*******************A****************G*****************************************A**G*******ACAG*AACA*GG*CC*TAG------------------
*FTF1a-3* ******************************G*******************A**C*********************AT**********************C*******************************************ACAG*AACA*GG*CA*TAG------------------
*FTF1c-1* ------------------------------------------------------------------------------------------------------------------------------------------------------------------------------------
*FTF1c-2* ------------------------------------------------------------------------------------------------------------------------------------------------------------------------------------
*FTF1c-3* ------------------------------------------------------------------------------------------------------------------------------------------------------------------------------------

Position 3961-3970
*FTF1a-1* TACCCGGTAG
*FTF1a-2* ----------
*FTF1a-3* ----------

FTF1c-1 ----------
FTF1c-2 ----------
FTF1c-3 ----------

**Supplementary Figure 24: Multiple sequence alignment of *FTF1* paralogs in MAFF 103036.** Alignment of nucleotide sequences comprising the complete ORF and the 500-bp upstream region of *FTF1a-1*, *FTF1a-2*, *FTF1a-3*, *FTF1c-1*, *FTF1c-2*, and *FTF1c-3*. Asterisks (*) indicate nucleotides identical to those of *FTF1a-1*, whereas nucleotide substitutions are explicitly shown. Gaps introduced to optimize the alignment are indicated by dashes (–). The translation start codon (ATG) of *FTF1a-1* is highlighted in red.
